# Smash++: an alignment-free and memory-efficient tool to find genomic rearrangements

**DOI:** 10.1101/2019.12.23.887349

**Authors:** Morteza Hosseini, Diogo Pratas, Burkhard Morgenstern, Armando J. Pinho

## Abstract

**Background:** The development of high-throughput sequencing technologies and, as its result, the production of huge volumes of genomic data, has accelerated biological and medical research and discovery. Study on genomic rearrangements is crucial due to their role in chromosomal evolution, genetic disorders and cancer;

**Results:** We present Smash++, an alignment-free and memory-efficient tool to find and visualize small- and large-scale genomic rearrangements between two DNA sequences. This computational solution extracts information contents of the two sequences, exploiting a data compression technique, in order for finding rearrangements. We also present Smash++ visualizer, a tool that allows the visualization of the detected rearrangements along with their self- and relative complexity, by generating an SVG (Scalable Vector Graphics) image;

**Conclusions:** Tested on several synthetic and real DNA sequences from bacteria, fungi, Aves and mammalia, the proposed tool was able to accurately find genomic rearrangements. The detected regions complied with previous studies which took alignment-based approaches or performed FISH (Fluorescence *in situ* hybridization) analysis. The maximum peak memory usage among all experiments was ~1 GB, which makes Smash++ feasible to run on present-day standard computers.

## 1 Background

With the ever-increasing development of high-throughput sequencing (HTS) technologies, a massive amount of genomic information is produced at much higher speed and lower cost than was possible before [1]. Analyses of such information has led to the advancement of our understanding of biology and disease, over the past decade [2, 3]. Computational solutions play a key role in dry-lab analysis of the deluge of HTS data by using efficient and fast algorithms.

Genome rearrangements are mutations that alter the arrangement of genes on a genome, and usually occur in the presence of errors in cell division following meiosis or mitosis. These structural abnormalities in chromosomes include, but are not limited to, deletions, duplications, translocations, inversions and insertions, mostly occur as an accident in the sperm or egg cell and hence are present in every cell of the body [4, 5].

Studies on chromosomal aberrations, which underlie many genetic diseases and cancer, are crucial for diagnostics, prognostics and targeted therapeutics [6, 7]. Examples of such diseases are the Wolf–Hirschhorn syndrome (WHS), that is caused by a partial deletion from human chromosome location 4p16.3 [8], the Charcot–Marie–Tooth disease (CMT), that is most commonly caused by duplication of the gene encoding peripheral myelin protein 22 (PMP22) on human chromosome 17 [9], and the acute myeloid leukemia (AML), that may be caused by translocations between human chromosome 8 and 21 [10].

In this paper, we present Smash++, an alignment-free tool that finds chromosomal rearrangements between two DNA sequences based on their information content, which is obtained by a data compression technique. This computational solution follows a combination of probabilistic and algorithmic approaches for having a quantitative definition of information, although it can be seen as more of a probabilistic one [11]. Associated with Smash++, we present a visualizer that is capable of visualizing as SVG images informationally similar regions between two genomic sequences. This tool also provides self- and relative redundancy (complexity) for the similar regions.

Smash++ is an improved version of Smash [12], featuring (1) improved accuracy, obtained by using multiple finite-context models along with substitution-tolerant Markov models to find fine-grained and coarse-grained chromosomal rearrangements, (2) presenting self-complexity (redundancy) and relative redundancy of informationally similar regions between two DNA sequences, (3) improved user interface (UI) in command line, by adding several options to customize the tool for running, and resulting SVG image, by adding markers for positions of DNA bases and also plotting self- and relative redundancy, and (4) improved performance, in terms of memory usage.

## 2 Results and Discussion

### 2.1 Implementation

Smash++ is implemented in the C++ language and is licensed under GNU GPLv3. It generates information maps for two sequences and, based upon that, finds similar regions in them, in which there can be potentially DNA rearrangements. Therefore, Smash++ gives an insight into positions of rearrangements that have happened between two sequences. The tool comes with a visualizer, that can be called in the command line with a flag called “-viz”. Similar regions in reference and target sequences are shown with the same color, that are chosen randomly using HSV color model. For more information about usage of the tool, see note S3 of the supplementary material.

The machine used for the tests had a 4-core 3.40 GHz Intel^®^ Core™ i7-6700 CPU and 32 GB of RAM. The Python script “xp.py”, in the “experiment” directory, can be used to reproduce the results by switching False/True the variables associated with each dataset.

### 2.2 Dataset

Smash++ and several other methods have been tested on a collection of synthetic and real sequences, that are described in Table 1. We used the GOOSE toolkit (https://github.com/pratas/goose) to make the synthetic sequences of which the sizes vary from 1.5 Kb to 100 Mb. We applied mutations and reversely complemented parts of the sequences. For a real dataset, we chose different sequences from bacteria, Aves, mammalia and fungi, with the sizes of ~1 Mb to ~ 127 Mb.

**Table 1:** Synthetic and real dataset used in the experiments. The real dataset can be download from NCBI via accession number (access.) provided in the descriptions.

| Sequence | Group | Length (b) | Description |
| --- | --- | --- | --- |
| GGA18 | Aves | 11,373,140 | Access.: CM000110 – *Gallus gallus* chromosome 18. |
| MGA20 | Aves | 10,730,484 | Access.: CM000981 – *Meleagris gallopavo* isolate NT-WF06-2002-E0010 breed Aviagen turkey brand Nicholas breeding stock chromosome 20. |
| GGA14 | Aves | 16,219,308 | Access.: CM000106 – *Gallus gallus* chromosome 14. |
| MGA16 | Aves | 14,878,991 | Access.: CM000977 – *Meleagris gallopavo* isolate NT-WF06-2002-E0010 breed Aviagen turkey brand Nicholas breeding stock chromosome 16. |
| HS12 | Mammalia | 133,275,309 | Access.: NC_000012 – *Homo sapiens* chromosome 12, GRCh38.p13 Primary Assembly. |
| PT12 | Mammalia | 130,995,916 | Access.: NC_036891 – *Pan troglodytes* isolate Yerkes chimp pedigree #C0471 (Clint) chromosome 12. |
| PXO99A | Bacteria | 5,238,555 | Access.: CP000967 – *Xanthomonas oryzae* pv. *oryzae* causes the major disease of bacterial blight of rice (*Oryza sativa* L.). *X. oryzae* pv. *oryzae* PXO99A strain is virulent toward a large number of rice varieties representing diverse genetic sources of resistance [21]. |
| MAFF 311018 | Bacteria | 4,940,217 | Access.: AP008229 – *X. oryzae* pv. *oryzae* MAFF 311018 is a Japanese race 1 strain [37]. |
| ScVII | Fungi | 1,090,940 | Access.: NC_001139 – *Saccharomyces cerevisiae* S288C chromosome VII. |
| SpVII | Fungi | 1,105,967 | Access.: CP020299 – *Saccharomyces paradoxus* strain UFRJ50816 chromosome VII. |
| RefS | Synthetic | 1,500 | It consists of three segments of 500 base size. |
| TarS | Synthetic | 1,500 | To build TarS, segment I is mutated 2%, II is inversely repeated and III is duplicated. |
| RefM | Synthetic | 100,000 | It has four segments of 25 kilobase size. |
| TarM | Synthetic | 100,000 | For building TarM, segment I of RefM (out of total four) is inversely repeated, II is mutated 90%, III is duplicated and IV is mutated 3%. |
| RefL | Synthetic | 5,000,000 | It includes two segments, 2,500,000 bases each. |
| TarL | Synthetic | 5,000,000 | Segment I is inversely repeated and II is mutated 2% for building TarL. |
| RefXL | Synthetic | 100,000,000 | It is made of four segments, 25,000,000 bases each. |
| TarXL | Synthetic | 100,000,000 | Segment I is mutated 1%, segments II and III are inversely repeated and segment IV is duplicated to make TarXL. |
| RefMut | Synthetic | 60,000 | It includes 60 segments of 1 kilobase size. |
| TarMut | Synthetic | 60,000 | To build TarMut, the first segment (I) is mutated 1%, the second segment is mutated 2%, the third one is mutated 3%, and so on. |
| RefComp | Synthetic | 1,000,000 | It consists of 10 segments of 100 kilobases. |
| TarComp | Synthetic | 1,000,000 | For building it, the first segment (I) of RefComp is duplicated, the second, third and fourth segments are mutated 1%, 2% and 3%, respectively. The segments V, VI and VII of RefComp are inversely repeated, then mutated 4%, 5% and 6%, respectively. Finally, the segments VIII, IX and X are mutated 7%, 8% and 9%, respectively. |
| RefPerm | Synthetic | 3,000,000 | It includes three segments of 1 megabase size. In addition to the original sequence, it is permuted, using GOOSE toolkit, by blocks of sizes 450 Kb, 30 Kb, 1 Kb and 30 b. |
| TarPerm | Synthetic | 3,000,000 | To build TarPerm, the first segment is mutated 1%, the second segment is inversely repeated and the third one is mutated 2%. |

### 2.3 Application on synthetic data

Figure 1 illustrates the result of running Smash++ and the associated visualizer on a synthetic dataset. The top sections show how we have built the reference and the target sequences. For example, to build the reference sequence in Figure 1a, we generated three random sequences of size 500 b, using GOOSE, and concatenated them. For building the target sequence, we made reverse complements of parts I and III from the reference, and also mutated part II 2%, then we concatenated the parts in the order shown in the figure. Figures 1b, c and d follow the same procedure. To build the target in Figure 1e, we mutated the first 1 Kb block of the reference 1%, the second block 2%, the third block 3%, up until the 60th block that we mutated 60%.

**Figure 1:**
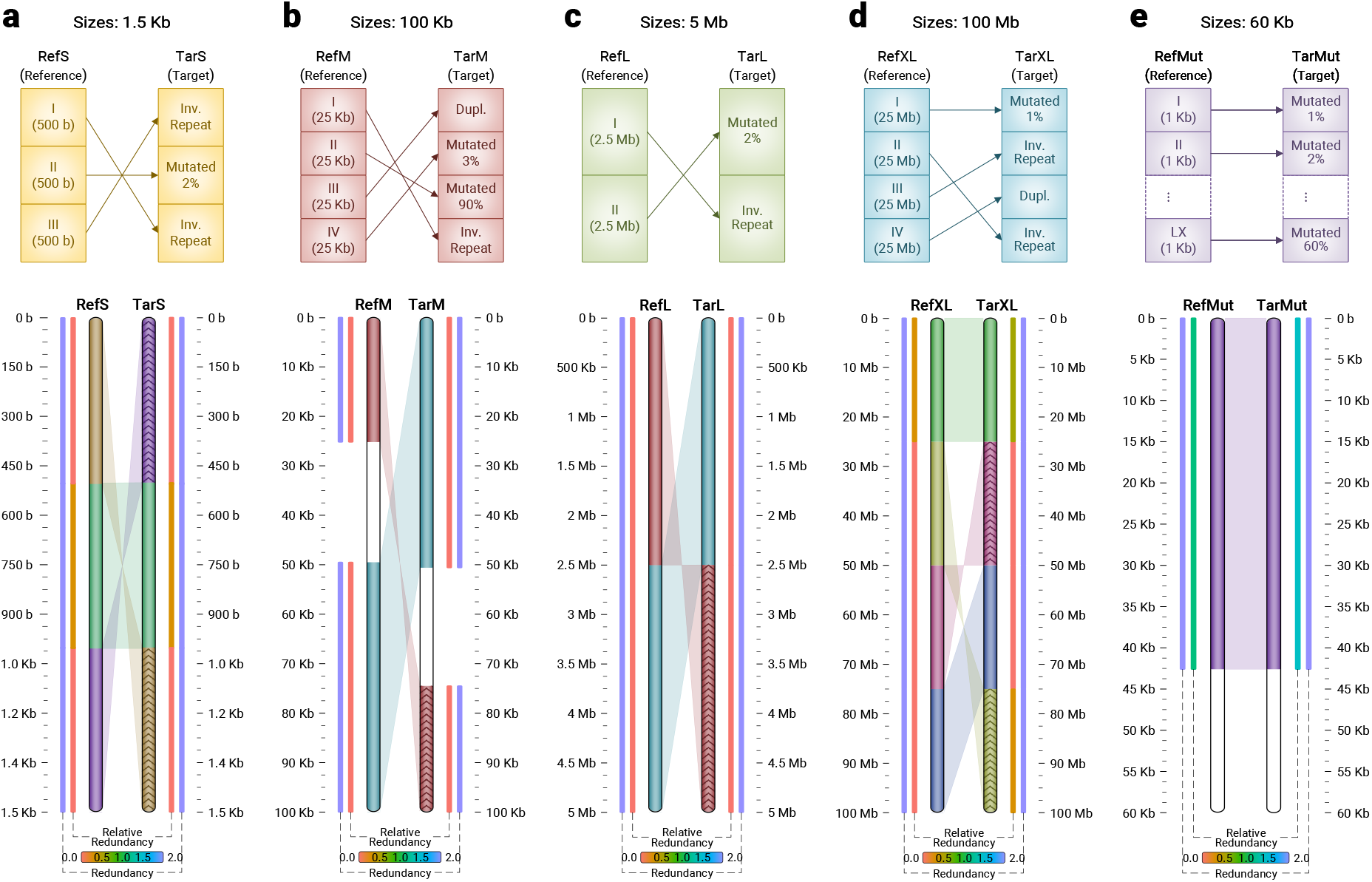
Similarities between synthetic sequences with different sizes, detected by Smash++. The parameters used are: *k*-mer size = 14 and number of substitutions in STMM = 5, which are the default parameters used by Smash++. For the threshold, the default value of 1.5 and 1.97 are used for subfigures a-d and e, respectively. (a) 1.5 Kb sequences; (b) 100 Kb sequences. No similarity is detected for part II of the reference, since it is mutated 90%. Parts III and IV of the reference and I and II of the target are joined, since there is no space between consecutive regions; (c) 5 Mb sequences; (d) 100 Mb sequences; (e) 60 Kb sequences. ~43% of mutation is detected.

The bottom sections of Figure 1 show the output of the Smash++ visualizer, detecting similar regions regardless of their sizes. Note that for each detected region, the average value of redundancy and relative redundancy is illustrated. In Figure 1b, part II of the reference is mutated 90%, i.e., nine out of every ten bases is mutated, on average. As expected, Smash++ does not recognize similarity between this pair of regions. Also, in the case of parts III and IV of the reference, since we detect similarity between part III of the reference and I of the target, and also part IV of the reference and II of the target, and there is no space between these regions, we join them and consider them as a bigger region of size 50 Kb. Figure 1e shows that Smash++ is able to detect ~43% of mutation, which has been made possible by the usage of substitution-tolerant Markov models (see section “Methods”). Figure 1 shows that Smash++ can be employed to detect small-scale and large-scale similarities between DNA sequences.

### 2.4 Application on real data

Figure 2 shows similarities between real sequences, found by Smash++. Subfigures a and b show similarities of chromosomes 18 and 14 of *Gallus gallus* (chicken) with orthologous chromosomes 20 and 16 of *Meleagris gallopavo* (turkey), respectively. These avian species, that are of great agricultural and commercial importance, are estimated to have diverged at 37.2 MYA [13]. Figures 2a and b demonstrate that Smash++ was able to find the inversions confirmed by FISH analysis, reported at [14, 15].

**Figure 2:**
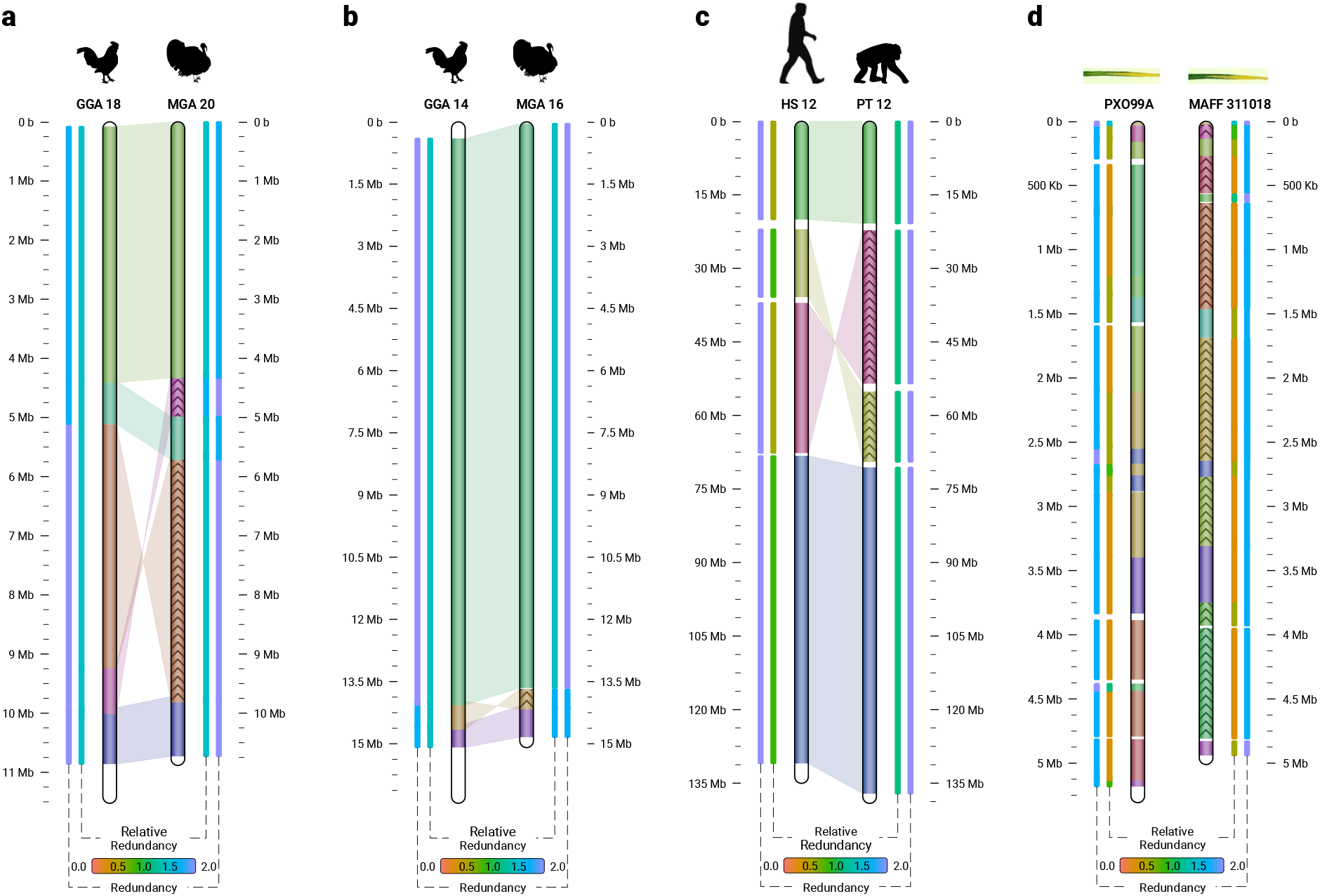
Similarities in a real dataset, detected by Smash++. (a) *G. gallus* (chicken) chr. 18 and *M. gallopavo* (turkey) chr. 20. The parameters were *k*-mer size = 14, no. substitutions in STMM = 5, threshold = 1.9 and min block size (*m*) = 500, 000, i.e., the regions smaller than 500,000 bases were not considered for further processing; (b) *G. gallus* chr. 14 and *M. gallopavo* chr. 16. The result is obtained by setting *k* = 14, no. substitutions = 5, threshold = 1.95 and *m* = 400, 000; (c) *H. sapiens* (human) chr. 12 and *P. troglodytes* (chimpanzee) chr. 12. The parameters were *k* = 14, without using STMM, threshold = 1.9 and *m* = 100, 000; (d) *X. oryzae* pv. *oryzae* PXO99A (a rice pathogen) and *X. oryzae* pv. *oryzae* MAFF 311018 (a rice pathogen). The result obtained by setting *k* = 13, threshold = 1.55 and m = 10, 000.

In Figures S1 and S2 of the supplementary material, we have compared Smash++ with other methods, on GGA 18 / MGA 20 and GGA 14 / MGA 16 chromosomes, respectively. The methods included in these figures are: (a) Smash++; (b) progressiveMauve [16], that uses an alignment objective score to detect rearrangement breakpoints when genomes have unequal gene content. It also applies a probabilistic alignment filtering method in order for removing erroneous alignments of unrelated sequences; (c) the method proposed in [15], that takes a bacterial artificial chromosome (BAC)-based approach along with FISH analysis to develop an integrated physical, genetic and comparative map of chicken and turkey; (d) SynBrowser [17], that constructs synteny blocks using prebuilt alignments in the UCSC genome browser database; and (e) FISH analysis [14].

Figure 2c demonstrates similarities between chromosomes 12 of *Homo sapiens* and *Pan troglodytes,* that are estimated to have diverged at 6.7 MYA. A comparison to other methods is provided in Figures S3 of the supplementary material. The methods include: (a) Smash++; (b) progressiveMauve; (c) Cinteny [18], that performs sensitivity analysis for synteny block detection and for the subsequent computation of reversal distances, by means of an extended version of ternary search trees (TST). Embedded in this extension are “walks” through the leaves of the tree, that correspond to walks on the genome markers in their linear order; (d) SynBrowser; and (e) D-Genies [19], that works based on alignment of genomes by minimap2 software package [20].

Figure 2d illustrates similarities between *Xanthomonas oryzae* pv. *oryzae* PXO99A and *Xanthomonas oryzae* pv. *oryzae* MAFF 311018, two strains of *Xanthomonas oryzae* pv. *oryzae* (Xoo) pathogen, which causes the disease of bacterial blight of rice *(Oryzae sativa* L.). That is the most serious bacterial disease of rice that can reduce yields by as much as 50% [21]. Note that to have a clearer picture, we have not plotted the shades connecting similar regions. This can be achieved by “-1 6” option while calling the Smash++ visualizer. Figure S4 of the supplementary material provides the comparison of Smash++ with progressiveMauve and the study [21], which uses an alignment method to find genome rearrangements in Xoo. As can be seen, the result provided by Smash++ conforms to the one presented in the study [21], without performing an alignment.

### 2.5 Comparison to Smash

To have a better understanding of the improvement we have made over the first version, Smash, we compare the two tools on a synthetic and a real dataset (see Figure 3). In Table 1, the procedure of making the synthetic data (RefComp and TarComp) is described. Figure 3a shows the comparison of running Smash and Smash++ on the synthetic dataset. For Smash, we used an FCM with *k*-mer size of 14, and for Smash++, we used a cooperation of an FCM with *k*-mer size of 14 and an STMM with number of substitutions of 5. As the information profiles show, Smash++ is able to model better the data, since it uses less information (lower information contents) to describe the target based on the reference; this is possible because of employing a cooperation of the FCM and the STMM instead of using solely an FCM. We expect the output to have the following format: parts I, II, III and IV of the reference and the target are similar (including rearrangements), there is also inverted repeats between parts V, VI and VII of the sequences, and finally, there are rearrangements between parts VIII, IX and X of the sequences. When there is no space between consecutive regions, Smash++ joins them; therefore, we expect Smash++ to detect three similar regions: the one including parts I, II, III and IV, the one with parts V, VI and VII, and the one including parts VIII, IX and X. The rearrangements map shows that Smash++ fulfills our expectation. On the other side, Smash was not able to detect all rearrangements, showing that to model such dataset, we need more than a single FCM.

**Figure 3:**
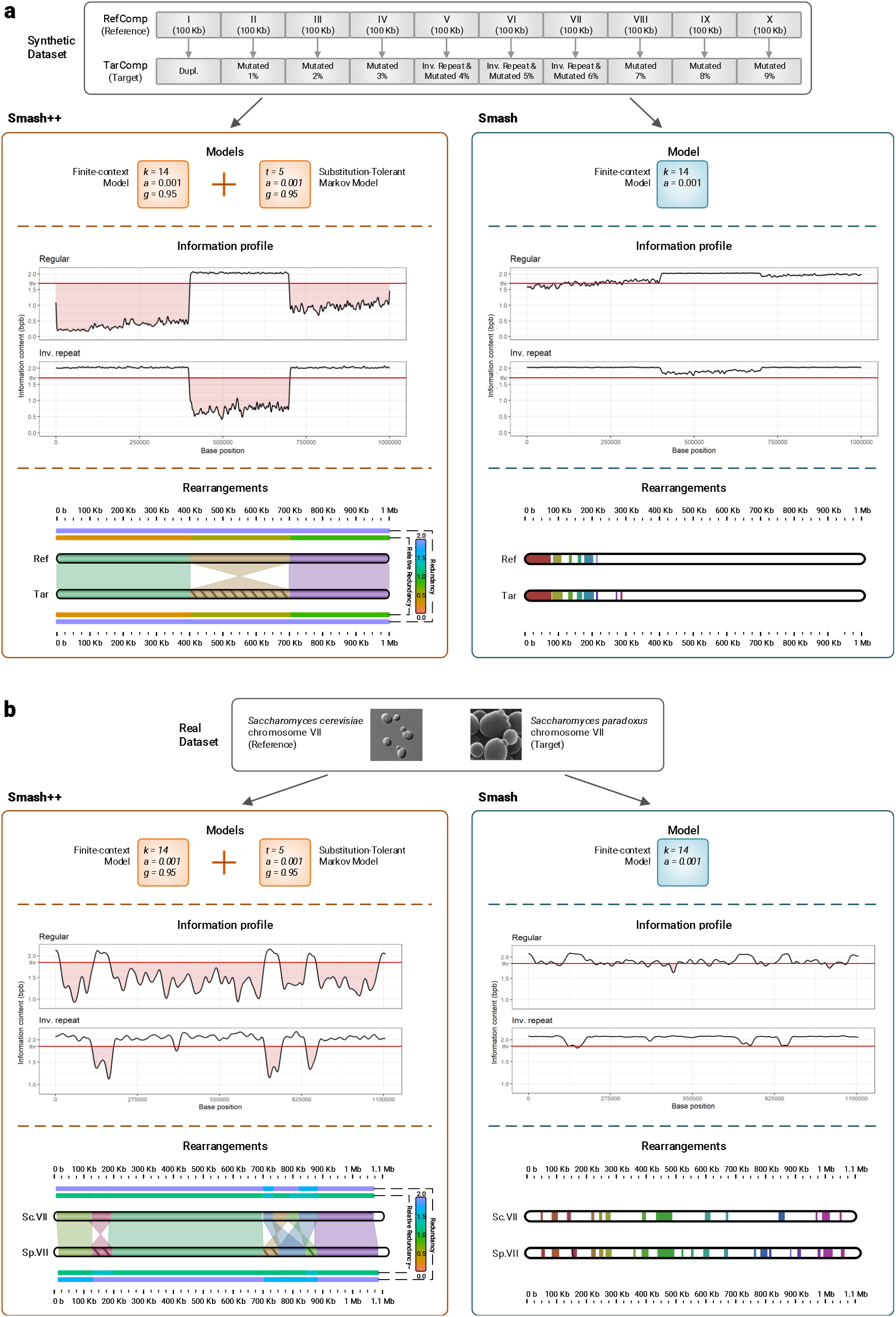
Comparison of Smash++ and Smash on (a) synthetic dataset. Using cooperation of an FCM and an STMM (in Smash++) produces more accurate results rather than using a single FCM (in Smash); and (b) real dataset, including *S. cerevisiae* chr. VII and *S. paradoxus* chr. VII. The rearrangements maps clearly show the improvement made over Smash, using an FCM along with an STMM.

The result of running Smash and Smash++ on a real dataset, *Saccharomyces cerevisiae* chromosome VII and *Saccharomyces paradoxus* chromosome VII, is demonstrated in Figure 3b. *S. cerevisiae* is a species of yeast that plays a key role in winemaking, baking and brewing. It has been a eukaryotic model organism that gives insights into molecular functioning of human cells [22]. *S. paradoxus* is closest known species to the *S. cerevisiae*, that has proved its importance on different fields of the life sciences, including evolution, ecology and biotechnology [23]. For the experiment, we ran Smash using an FCM with *k*-mer size of 14, and Smash++ using an FCM with *k*-mer size of 14 cooperated with an STMM with number of substitutions of 5. As can be seen, using an FCM along with an STMM could drastically improve modeling the data, which led to find rearrangements more accurately. The rearrangements map of Smash++ conforms to the previous study [22].

### 2.6 Robustness against fragmented data

Inherited from Smash, Smash++ is capable of finding similarities between a fragmented reference and a target sequence. Figure S5 of the supplementary material shows robustness of the proposed tool against fragmented data, for different randomly permutated block sizes. As can be seen, the same three target regions are detected even when the reference is fragmented to 100,000 blocks of 30 bases. This capability might be of interest in case of non-assembled sequences or in presence of assembly errors; note that this approach can not be considered as an alternative to assembly.

### 2.7 Benchmarking

Figure 4 illustrates performance of the proposed tool in terms of memory and time usage for all datasets (for more details, see supplementary Table S1). Size of the datasets are mentioned on top of each circle, in black, and number of detected similar regions between each pair of sequences is mentioned inside the circles, in white. The legend shows the precise size of datasets in bases (b). Figure 4a shows the peak memory in gigabytes used by Smash++ on all synthetic and real datasets. The maximum peak memory usage, ~1.08 GB, was when the proposed tool was ran on human and chimpanzee chromosomes 12, that are the biggest datasets with the total size of ~268 Mb. It should be mentioned that the memory usage of Smash++ is related to the *k*-mer size that is used for modeling the data, since different data structures are employed for different *k*-mer sizes (see section “Methods”). The sizes 13 and 14 were used to perform the experiments. The maximum memory usage of ~1 GB enables Smash++ to run on any computer, nowadays.

**Figure 4:**
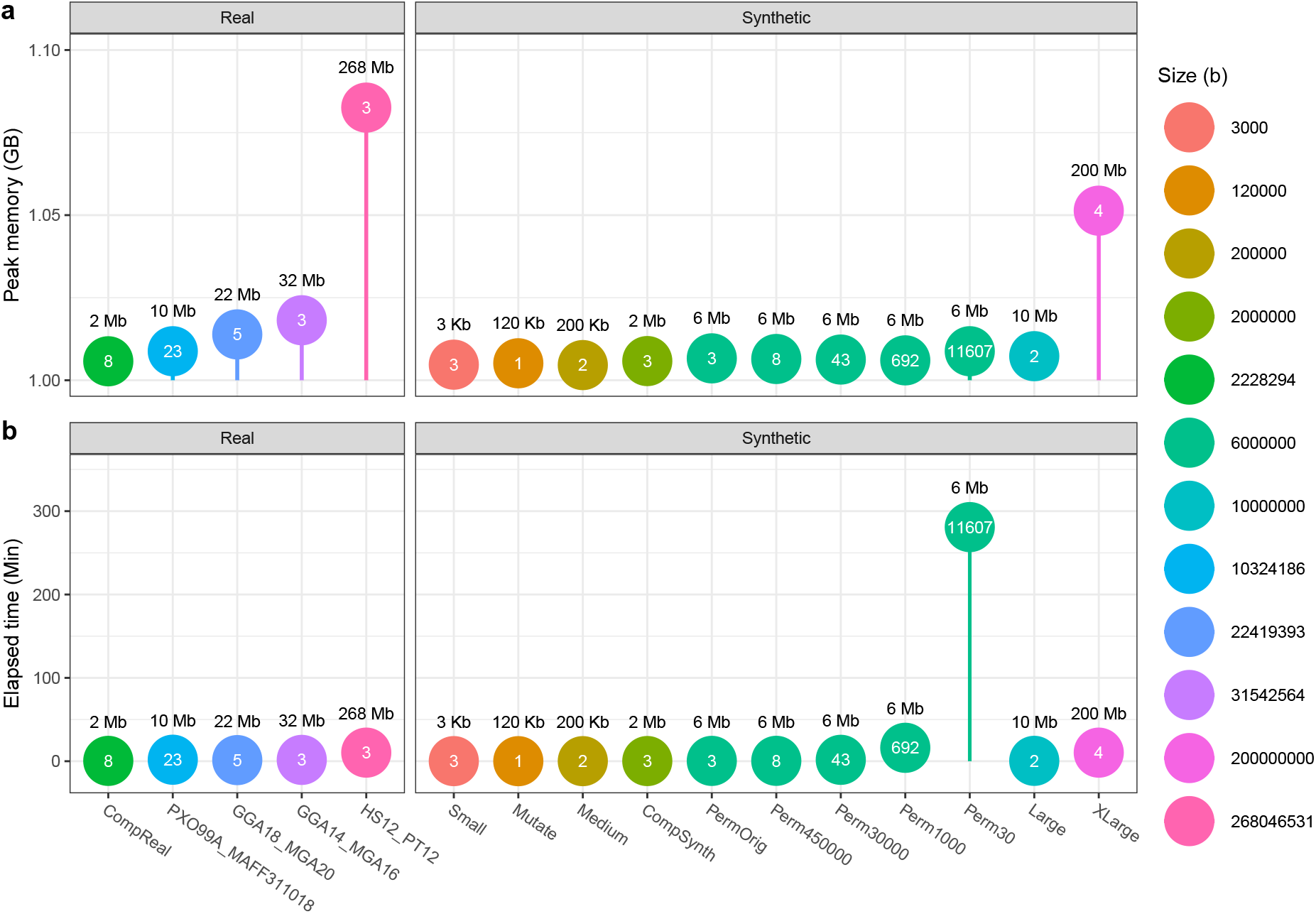
The peak memory consumption, in gigabytes, and elapsed (wall clock) time usage, in minutes, of Smash++ obtained by carrying out on all synthetic and real datasets described in Table 1.

Figure 4b demonstrates elapsed (wall clock) times, in minutes. The elapsed times rely on the file sizes along with the number of detected similar regions, meaning that the more the number of regions is and/or the greater the dataset size is, the more the time will be taken. Note that it is not a linear relation. As an example, the pair datasets “Large” and the pair “PXO99A_MAFF311018” have approximately the same total size of 10 Mb. In the former case that two similarities is detected, Smash++ takes ~26 seconds, but in the latter case with 23 similarities, the proposed tool takes ~1.8 minutes to run. As another example, carrying out Smash++ on the pair “XLarge” with the total size of 200 Mb and four similarities detected, takes approximately the same wall clock time as carrying out on the pair “HS12_PT12” that has the total size of ~268 Mb and three similar regions are detected. Regarding the pair “Perm30” with 11,607 similarities detected, we should notice that it has a massively fragmented reference sequence with 10,000 fragments of 30 b, therefore it is by far the most time-consuming dataset. Note that the difference between the values of 10,000 (number of reference fragments) and 11,607 (number of similar regions) arises from the fact that a number of the reference chunks are similar to more than one target region and vice versa. It is worth mentioning that due to the absence of a tool that provides relative and self-complexity in addition to detecting similarities, we cannot have a fair camparison to other tools in terms of time and memory usage; therefore, we have only provided the performance results for the proposed tool.

## 3 Conclusions

Finding genomic rearrangements is crucial, since they play an important role in genetic disorders, cancer and chromosomal evolution. We presented Smash++, an alignment-free tool that accurately finds small- and large-scale genomic rearrangements between pairs of DNA sequences, by employing a data compression approach. This memory-efficient tool was successfully tested on several synthetic and real data from bacteria, fungi, Aves and mammalia. The presented results showed that the detected rearrangements complied with previous studies which used alignment-based methods or performed FISH analysis. Smash++ consumed a maximum of ~1 GB of memory, among all experiments, which showed that it can be run on any computer, nowadays. The proposed tool has the potential to improve accuracy of diagnostic and genetic counselling, and also to guide future inverstigations into development of personalized therapeutic.

## 4 Methods

The schema of the proposed method is illustrated in Figure 5. Smash++ takes as inputs a reference and a target sequence and produces as output a position file, including local similarities of the two sequences, which can then be used by the Smash++ visualizer to produce an SVG image illustrating the similarities. This process has eight major stages: (1) compression of the original target file, based on the model of the original reference file, (2) filtering the information profile, which is the output of stage 1, and segmenting the target sequence, (3) reference-free compression of the segmented sequences, obtained by the previous stage, (4) compression of the original reference file, based on the model of segmented sequences, which are obtained by stage 2, (5) filtering the information profile and segmenting the reference sequence, (6) reference-free compression of the segmented sequences, (7) aggregating positions, that are generated by stages 3 and 6, and (8) visualizing the positions. The following sections describe the process in detail.

**Figure 5:**
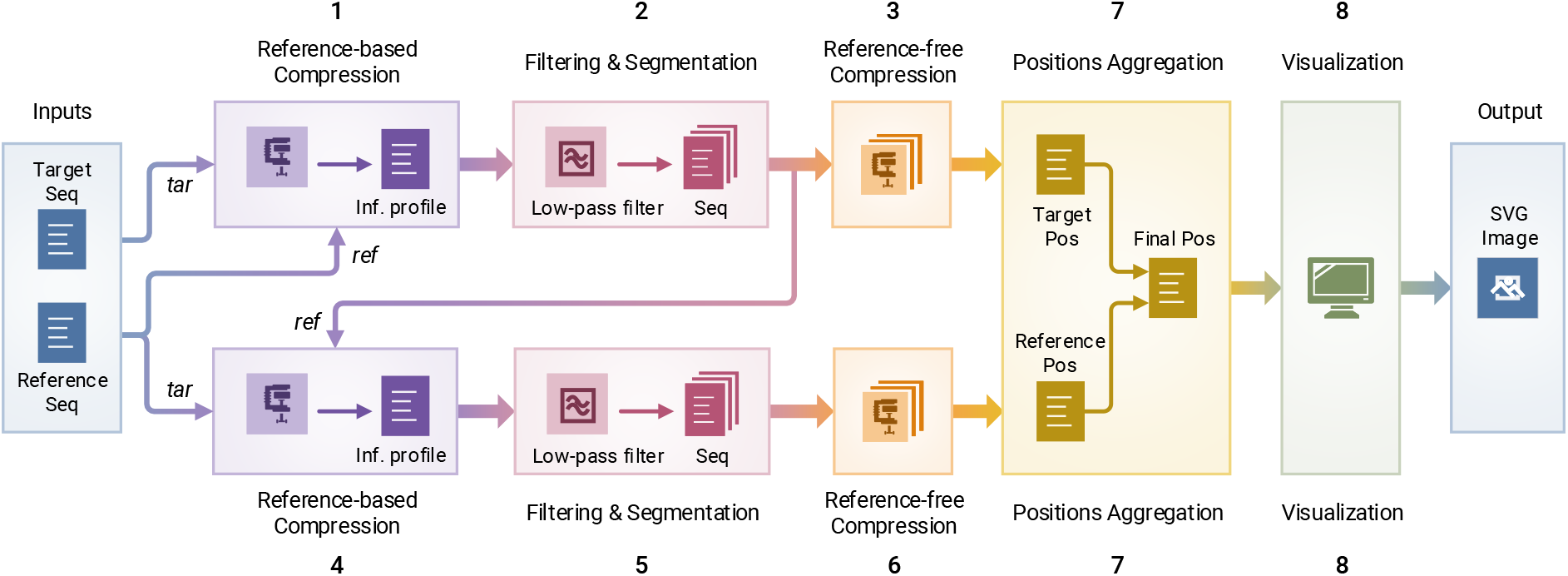
The schema of Smash++. The process of finding similar regions in reference and target sequences. Computing the redundancy in each region includes eight stages. Finally, Smash++ outputs a *.pos file that includes the positions of the similar regions, and can be then visualized, resulting in an SVG image.

### 4.1 Data modeling

We consider sequences over the nucleotide alphabet Θ = {A, C, G, T}; our goal is to measure the degree of local similarity between two such sequences. More specifically, we consider a reference sequence *S* = *s*_1_,…, *s_N_* over Θ, and we want to measure the local *information content* of a target sequence, given this reference sequence. To this end, we employ a combination of finite-context models and substitution-tolerant Markov models to derive different probablitly measures for observing a nucleotide *x* in a sequence, given the *context* of the previous *k* nucleotides (Fig. 6a); these probabilities are then mixed (by multiplications and additions shown in Fig. 6b) to provide the final probability (*P*) of observing the nucleotide *x*. The following subsections describe the models we use in detail.

**Figure 6:**
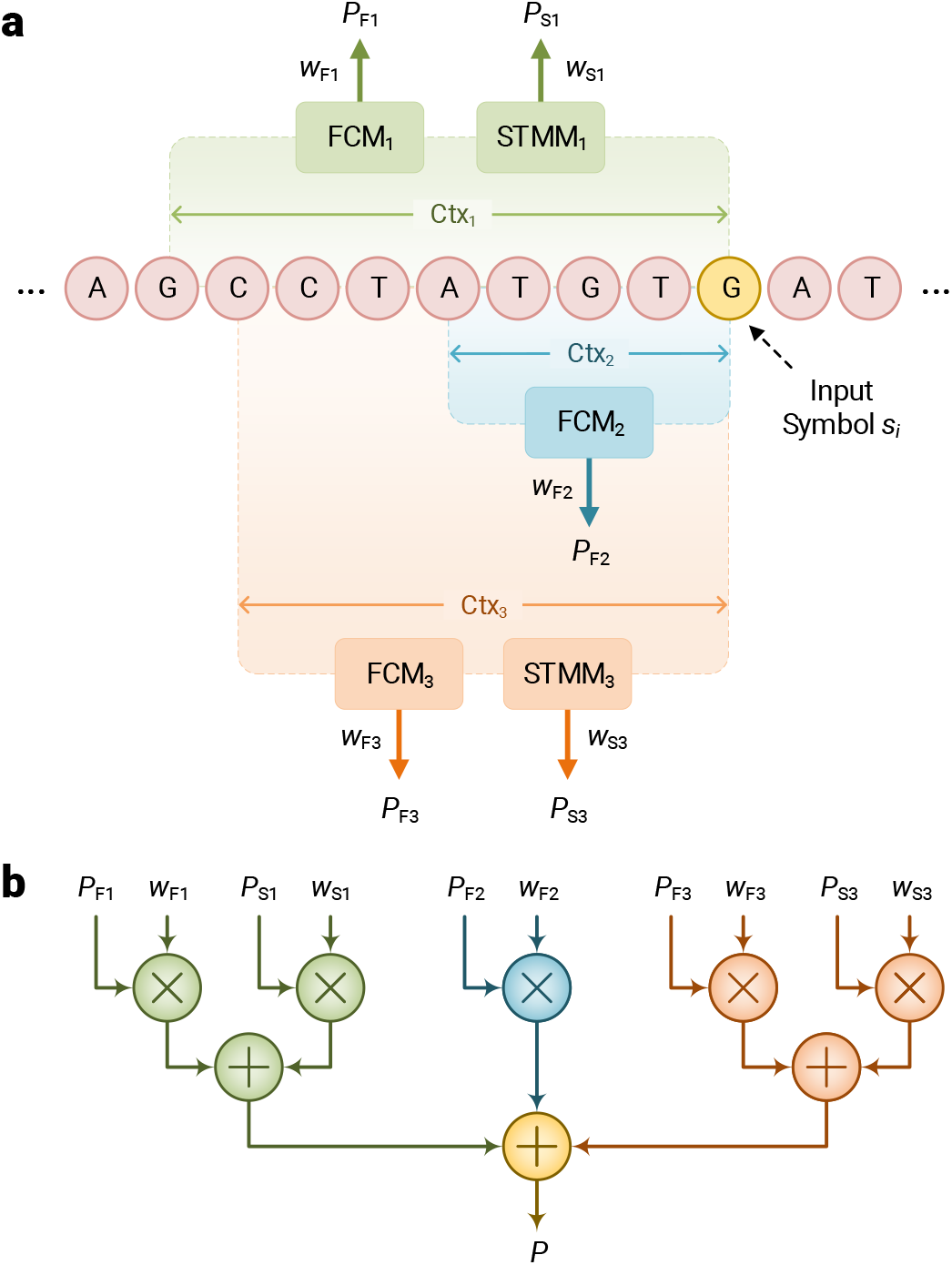
Data model used by Smash++. (a) cooperation between finite-context models (FCMs) and substitution-tolerant Markov models (STMMs). Note that each STMM needs to be associated with an FCM. (b) probability of an input symbol is estimated by employing the probability and weight values that have been obtained from processing previous symbols.

#### Finite-context model (FCM)

We consider the probability of observing a certain nucleotide, given the previous *k* nucleotides, by using the relative frequency of this event in the reference sequence *S*. For *x* ∈ Θ and a *k*-mer *Q* ∈ Θ^*k*^, let *N*(*x*|*Q*) be the number of occurrences of *Q* in *S* that are followed by nucleotide *x*, and let *N*(*Q*) be the number of occurrences of *Q* in *S*. As in [11, 24, 25], we then define

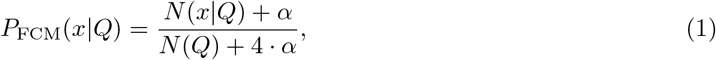

where “4” is the size of alphabet Θ and *α* is a pseudo-count parameter. For *α* =1, Eq. 1 turns into the *Laplace estimator*. Note that an FCM has the Markov property, in which the conditional probability distribution of observing a nucleotide depends only upon the state of preceding *k*-mer.

#### Substitution-tolerant Markov model (STMM)

Given the reference sequence *S*, we use the aforementioned probability distribution *P*_FCM_ to define a sequence 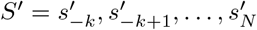 recursively by

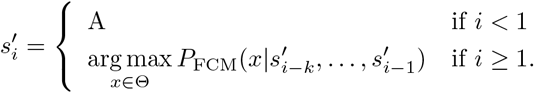

For *x* ∈ Θ and a *k*-mer *Q* ∈ Θ^*k*^, we then define *N*’(*x*|*Q*) as the number of occurrences of *Q* followed by *x* and *N*’(*Q*) as the number of occurrences of *Q*, respectively, in the sequence *S*’. Finally, we define

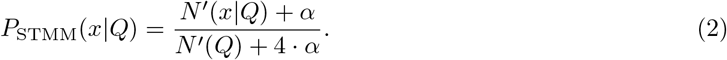

STMMs, that are probabilistic-algorithmic models [26, 11], can be used along with FCMs to modify the behavior of Smash++ when confronted with nucleotide substitutions in genomic sequences. These models can be disabled, to reduce the number of mathematical calculations, and consequently, increase the performance of the proposed method. Such operation is automatically performed using an array of size *k* (the context size), named history, which preserves the past *k* hits/misses. Observing a symbol in the sequence, the memory is checked for the symbol with the highest number of occurrences. If they are equal, a hit is saved in the history array; otherwise, a miss is inserted into the array. Before getting to store a hit/miss in the array, it is checked for the number of misses and in the case they are more than a predefined threshold *t*, the STMM will be disabled and also the history array will be reset. This process is performed for each nucleotide in the sequence.

The following example shows the distinction between an FCM and an STMM. Assume that the current context at a certain position is AGACGTAC, and the number of occurrences of symbols saved in memory is 10, 6, 15 and 8 for A, C, G and Ts, respectively; also, the symbol to appear in the sequence is T. An FCM considers the next context as GACGTAC**T**, while an STMM considers it as GACGTAC**G**, since the nucleotide G is the most probable symbol, based on the number of occurrences stored in memory.

#### Cooperation of FCMs and STMMs

When FCMs and STMMs are in cooperation, the probability of observing a nucleotide *x* in a sequence *S* can be estimated as

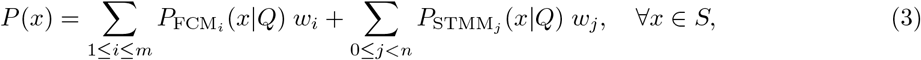

in which *m* and *n* denote the number of FCMs and STMMs, respectively, and *w_i_* and *W_j_* are weights assigned to each FCM and STMM, respectively, based on its performance. We have

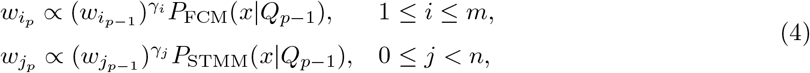

where *p* denotes a certain position, and *γ_i_* and *γ_j_* ∈ [0,1) are forgetting factors predefined for each model. Also,

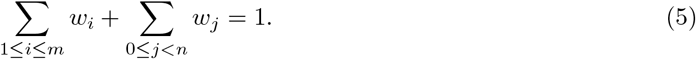

By experimenting different forgetting factors for models, we have found that higher factors should be assigned to models that have higher context-order sizes (less complexity) and vice versa. As an example, when the context size *k* = 6, *γ_i_* or γ_*j*_ ≃ 0.9 and when *k* = 18, *γ_i_* or *γ_j_* ≃ 0.95 would be appropriate choices. These values show that forgetting factor and complexity of a model are inversely related.

### 4.2 Storing models in memory

The FCMs and STMMs include, in fact, count values which need to be saved in memory. For this purpose, four different data structures have been employed considering the context-order size *k*, as follows:

- table of 64 bit counters, for 1 ≤ *k* ≤ 11,
- table of 32 bit counters, for *k* = 12,13,
- table of 8 bit approximate counters, for *k* = 14, and
- Count-Min-Log sketch of 4 bit counters, for *k* ≥ 15.

The table of 64 bit counters, that is shown in Figure 7a, simply saves the number of events for each context. The table of 32 bit counters saves in each position the number of times that the associated context is observed. When a counter reaches the maximum value 2^32^ — 1 = 4294967295, all the counts will be renormalized by dividing by two, as shown in Figure 7b.

**Figure 7:**
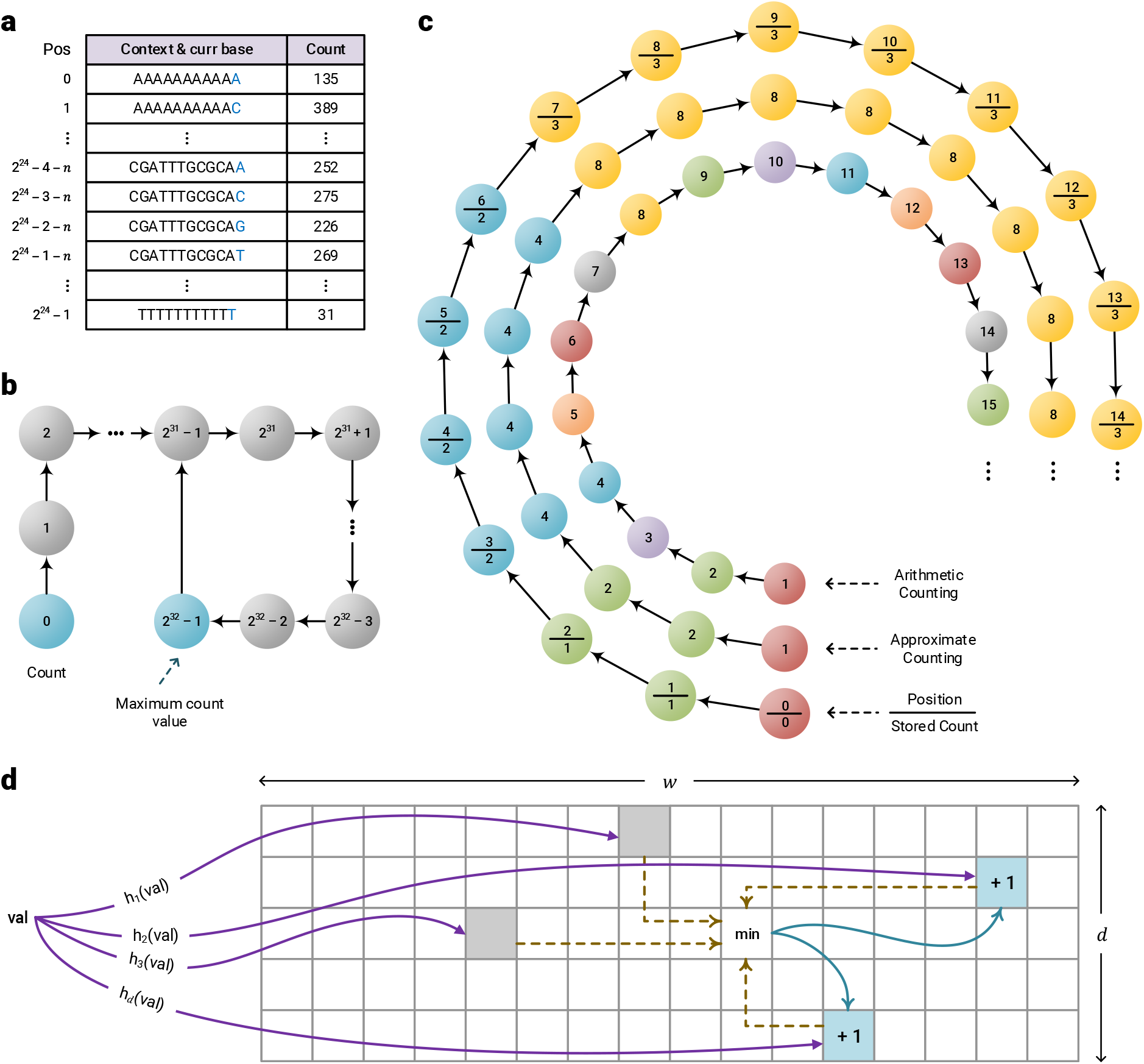
The data structures used by Smash++ to store the models in memory. (a) table of 64 bit counters that uses up to 128 MB of memory, (b) table of 32 bit counters that consumes at most 960 MB of memory, (c) table of 8 bit approximate counters with memory usage of up to 1 GB and (d) Count-Min-Log sketch of 4 bit counters which consumes up to 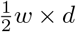 B of memory, e.g., if *w* = 2^30^ and *d* = 4, it uses 2 GB of memory.

Approximate counting is a method that employs probabilistic techniques to count large number of events, while using a small amount of memory [27]. Figure 8 shows the algorithm for two major functions associated with this method, U_PDATE_ and Q_UERY_. In order to update the counter, a pseudo-random number generator (PRNG) is used the number of times of the counter’s current value to simulate flipping a coin. If it comes up 0/Heads each time or 1/Tails each time, the counter will be incremented. Figure 7c shows the difference between arithmetic and approximate counting, and also the values which are actually stored in memory. Note that since an approximate counter represents the actual count by an order of magnitude estimate, one only needs to save the exponent. For example, if the actual count is 8, we store in memory log_2_ 8 = 3.

**Figure 8:**
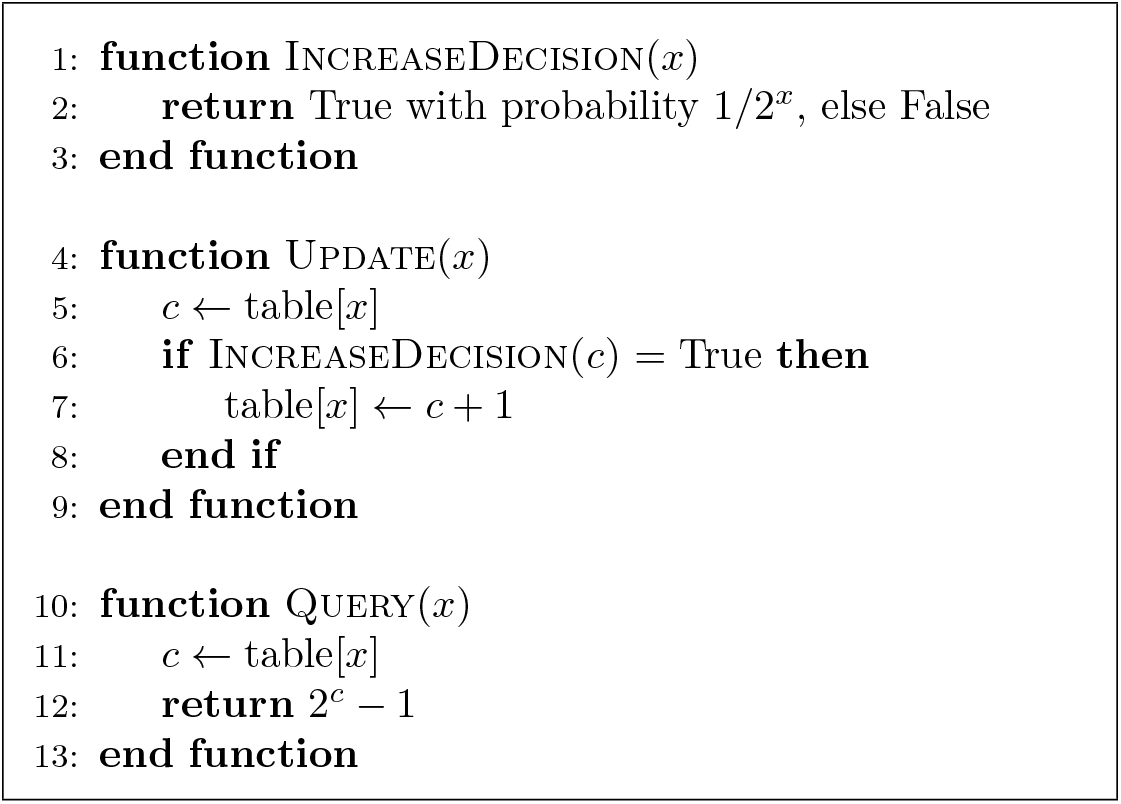
Approximate counting update and query.

Count-Min-Log Sketch (CMLS) is a probabilistic data structure to save frequency of events in a table by means of a family of independent hash functions [28]. The algorithm for updating and querying the counter is shown in Figure 9. In order to update the counter, its current value is hashed with *d* independent hash functions. Then, a coin is flipped the number of times of the counter’s current value, employing a pseudo-random number generator. If it comes up 0/Heads each time or 1/Tails each time, the minimum hashed values (out of *d* values) will be updated, as shown in Figure 7d.

**Figure 9:**
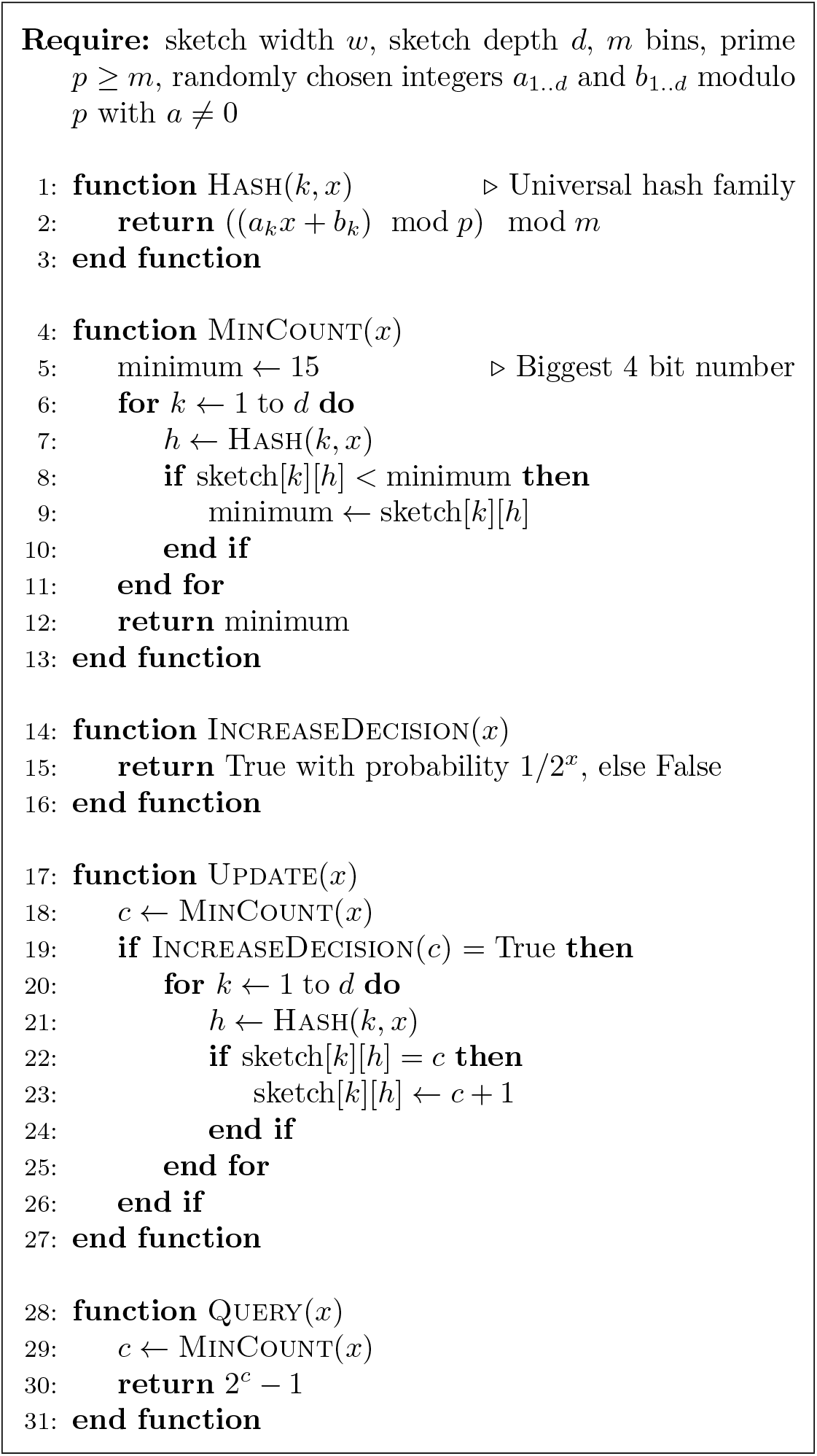
Count-Min-Log Sketch update and query.

CMLS requires a family of pairwise independent hash functions *H* = {*h: U* → [*m*]}, in which each function *h* maps some universe *U* to *m* bins. In this family of functions, the probability that all *x,y* ∈ *U, x* = *y* will hash to any pair of hashed values *z*_1_, *z*_2_ is as if they were perfectly random, i.e., *P*_*h*∈*H*_ [*h*(*x*) = *z*_1_ Λ *h*(*y*) = *z*_2_] = 1/*m*^2^. A hash function in this family can be obtained by

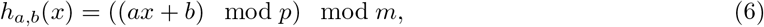

where *p* ≥ *m* is a prime number and *a* and *b* are randomly chosen integers modulo *p* with *a* ≠ 0. Note that if the number of bins is a power of two, *m* = 2^*M*^, multiply-add-shift scheme [29] can be used to avoid modular arithmetic. A hash function in this scheme can be obtained by:

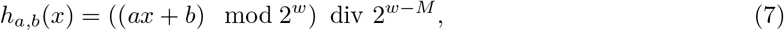

in which *w* is the number of bits in a machine word, e.g., 64, *a* is a random positive integer less than 2^*w*^ and *b* is a random non-negative integer less than 2^*w–M*^. Such hash function can be implemented in the C++ language by

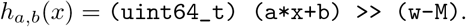

### 4.3 Finding similar regions

To find similar regions in reference and target sequences, a quantity is required for measuring the similarity. We use “per symbol information content”, in bpb (bit per base), which can be calculated as

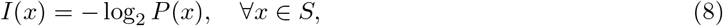

where *P*(*x*) denotes the probability of observing a nucleotide *x* in the sequence *S*, obtained by Equation 3.

The information content is the amount of information required to represent a symbol in the target sequence, based on the model of the reference sequence. The less the value of this measure is for two regions, the more amount of information is shared between them, and, therefore, the more similar are the two regions. Note that a version of this measure has been introduced in [12], which employs a single FCM to calculate the probabilities. In this paper, however, we exploit a cooperation between multiple FCMs and STMMs for highly accurate calculation of such probabilities.

The procedure of finding similar regions in a reference and a target sequence, illustrated in Figure 10, is as follows: after creating the model of the reference, the target is compressed based on that model and the information content is calculated for each symbol in the target. Then, the content of the whole target sequence is smoothed by a Hann window [30], which is a discrete window function given by 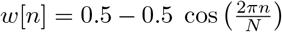, where 0 ≤ *n* ≤ *N* and length of the window is *N* +1. Next, the smoothened information content is segmented considering a predefined threshold, meaning that the regions with the content greater than the threshold are filtered out. This is carried out for both regular and inverted repeat homologies and, at the end, the result would be the regions in the target sequence that are similar to the reference sequence (Figure 10a). The described phase repeats for all of the target regions found, in the way that after creating the model for each region, the whole reference sequence is compressed to find those regions in the reference that are similar to each of the target regions (Figure 10b). The final result would have the form of Figure 10c.

**Figure 10:**
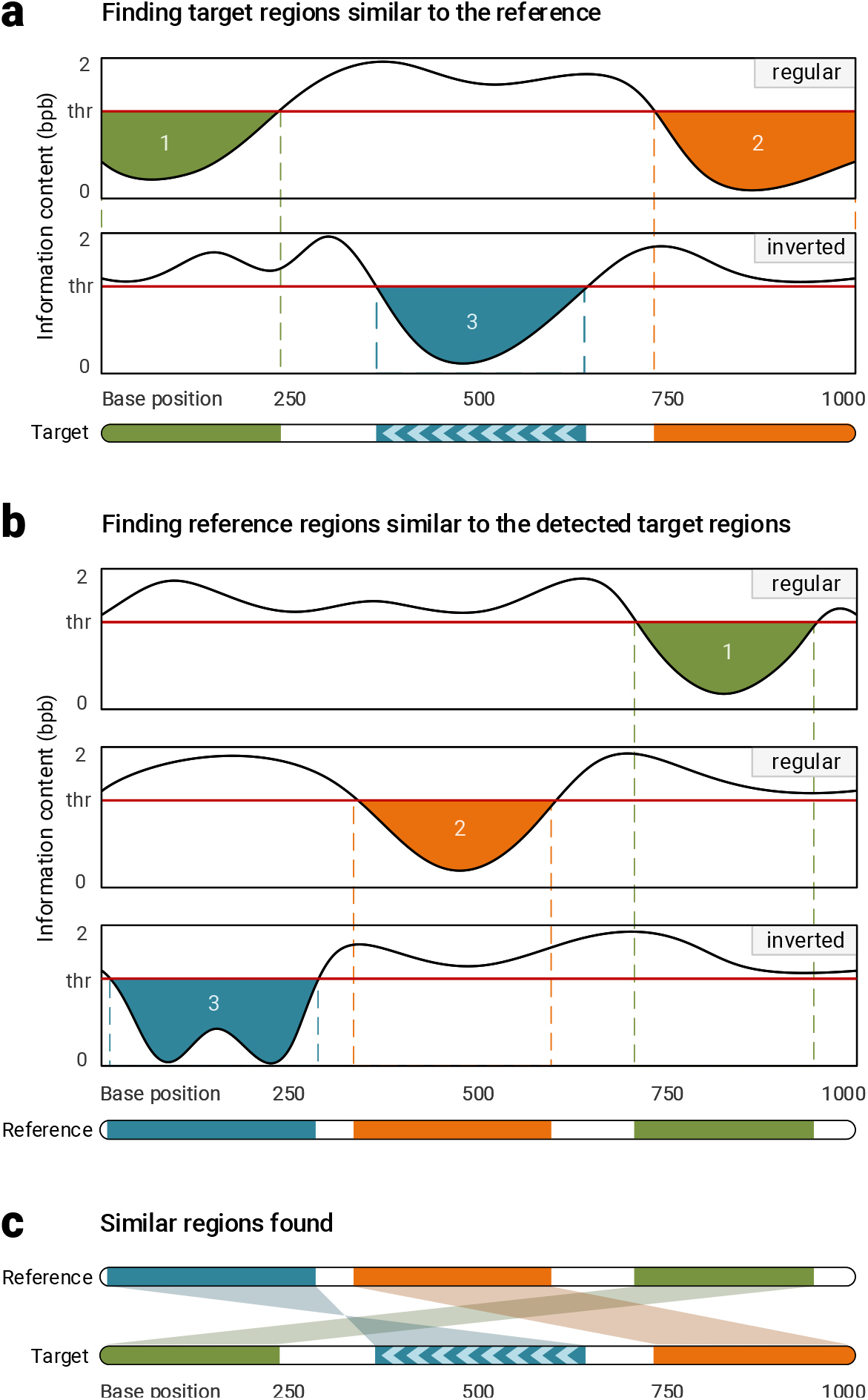
Finding similar regions in reference and target sequences. Smash++ finds, first, the regions in the target that are similar to the reference, and then, finds the regions in the reference that are similar to the detected target regions. This procedure is performance for both regular and inverted homologies.

### 4.4 Computing complexity

After finding the similar regions in reference and target sequences, we evaluate redundancy in each region, knowing that it is inversely related to Kolmogorov complexity, i.e., the more complex a sequence is, the less redundant it will be [31]. The Kolmogorov complexity, *K*, of a binary string *s*, of finite length, is the length of the smallest binary program *p* that computes *s* in a universal Turing machine and halts. In other words, *K*(*s*) = |*p*| is the minimum number of bits required to computationally retrieve the string *s* [32, 33].

The Kolmogorov complexity is not computable, hence, an alternative is required to compute it approximately. It has been shown in the literature that a compression algorithm can be employed for this purpose [34, 35, 36]. In this paper, we employ a reference-free compressor to approximate the complexity and, consequently, the redundancy of the found similar regions in the reference and the target sequences. This compressor works based on cooperation of FCMs and STMMs, which has been previously described in detail. Note that the difference between reference-based and reference-free version of such compressor is that, in the former mode, a model is first created for the reference sequence and, then, the target sequence is compressed based on that model, while in the latter mode, the model is progressively created at the time of compressing the target sequence.

## Availability of source code and requirements

- Project name: Smash++
- Project home page: https://github.com/smortezah/smashpp
- Operating system(s): Linux, macOS, Windows
- Programming language: C++, Python
- Other requirements: C++ 14, Python 3
- License: GNU GPLv3

## Availability of supporting data and materials

The data sets supporting the results of this article are available in the Smash++ Github repository, https://github.com/smortezah/smashpp/experiment/dataset.

## Additional files

A single Supplementary notes file, including:

**Supplementary Figure S1**. Comparison of Smash++ and other methods on *G. gallus* chromosome 18 and *M. gallopavo* chromosome 18.

**Supplementary Figure S2**. Different methods ruuning on *G. gallus* chromosome 14 and *M. gallopavo* chromosome 16.

**Supplementary Figure S3**. Comparing with other methods on *H. sapiens* chromosome 12 and *P. troglodytes* chromosome 12.

**Supplementary Figure S4**. Result of running different methods on *X. oryzae* pv. *oryzae* PXO99A and *X. oryzae* pv. *oryzae* MAFF 311018.

**Supplementary Figure S5**. Similarity of a target sequence to a fragmented reference sequence, that is randomly permutated by different block sizes.

**Supplementary Table S1**. Performance of Smash++ running on all synthetic and real datasets.

**Supplementary Note S1**. Software manual for Smash++.

## Declarations

## List of abbreviations

AML: acute myeloid leukemia;
BAC: bacterial artificial chromosome;
CMT: Charcot–Marie–Tooth;
CMLS: Count-Min-Log Sketch;
CPU: central processing unit;
FCM: finite-context model;
FISH: Fluorescence *in situ* hybridization;
GB: gigabyte;
GGA: *Gallus gallus*;
GHz: gigahertz;
HS: *Homo sapiens*;
HSV: Hue, Saturation, Value;
HTS: high-throughput sequencing;
KB: kilobyte;
MB: megabyte;
MGA: *Meleagris gallopavo;*
MYA: million years ago;
NCBI: national center for biotechnology information;
PMP22: peripheral myelin protein 22;
PRNG: pseudo-random number generator;
PT: *Pan troglodytes*;
RAM: random access memory;
Sc: *Saccharomyces cerevisiae*;
Sp: *Saccharomyces paradoxus*;
STMM: substitution-tolerant Markov model;
SVG: Scalable Vector Graphics;
TST: ternary search tree;
UI: user interface;
WHS: Wolf–Hirschhorn syndrome;
UCSC: University of California, Santa Cruz.

## Ethical Approval

Not applicable.

## Consent for publication

Not applicable.

## Author’s Contributions

M.H. developed the software and wrote the manuscript. D.P. and A.J.P. contributed to and tested the software. D.P., B.M. and A.J.P. provided guidance. All authors contributed to the manuscript.

## Acknowledgements

We would like to thank everyone who has contributed to the development of Smash++, through testing and feedback.

