## Supplementary for "Smash++: an alignment-free and memory-efficient tool to find genomic rearrangements"

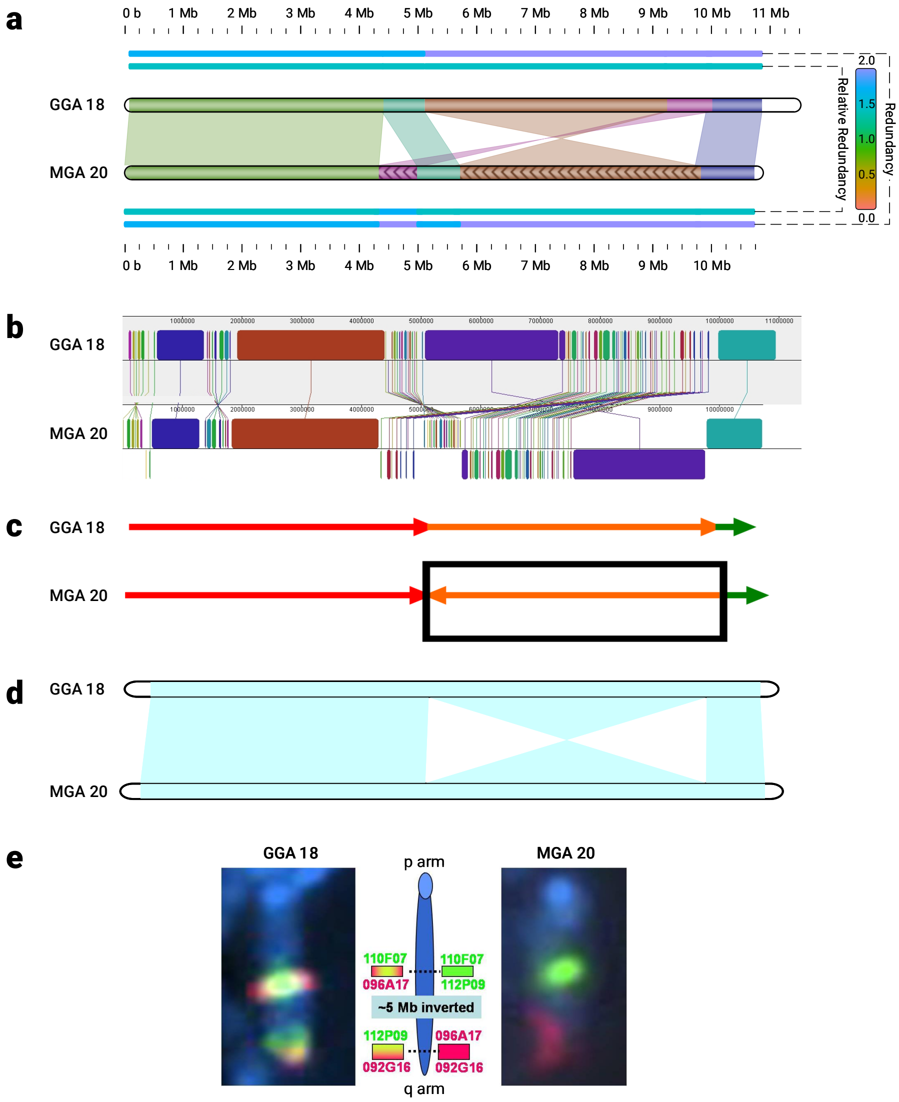


**Figure S1.** Pair-wise comparison of *G. gallus* chromosome 18 and *M. gallopavo* chromosome 20. (a) Smash++, with $k = 14$ and 5 used by an FCM and an STMM, respectively. The blocks smaller than 500 Kb are discarded; (b) progressiveMauve [1], with LCB (locally collinear block) weight of 18692. Reverse complements are shown in lower level; (c) adopted from [2], which is confirmed by fluorescence *in situ hybridization* (FISH) analysis. The box shows a local rearrangement; (d) SynBrowser [3], with the resolution of 150 Kb (minimum size of a reference block); (e) adopted from [4], which confirms an inversion rearrangement of size ∼5 Mb by FISH analysis.

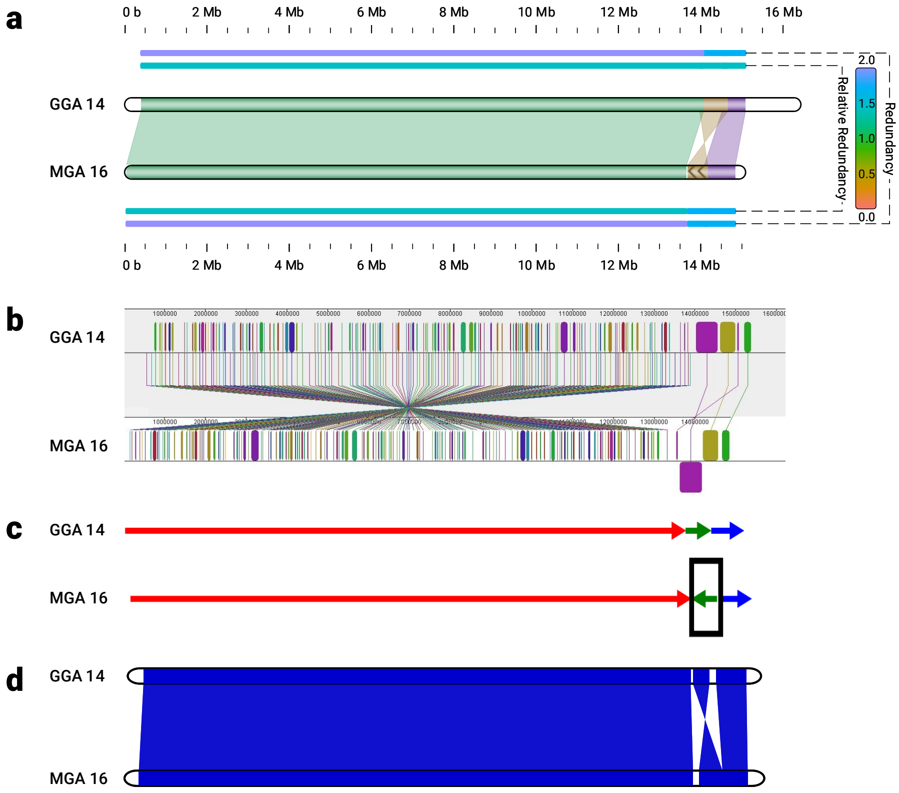


**Figure S2.** *G. gallus* chromosome 14 compared to *M. gallopavo* chromosome 16. (a) Smash++, employing an FCM and an STMM with $k = 14$ and 5, respectively. The blocks smaller than 400 Kb are discarded; (b) progressiveMauve, with LCB weight of 27424; (c) adopted from [2]. The box shows an inversion rearrangement; (d) SynBrowser, with the resolution of 150 Kb.

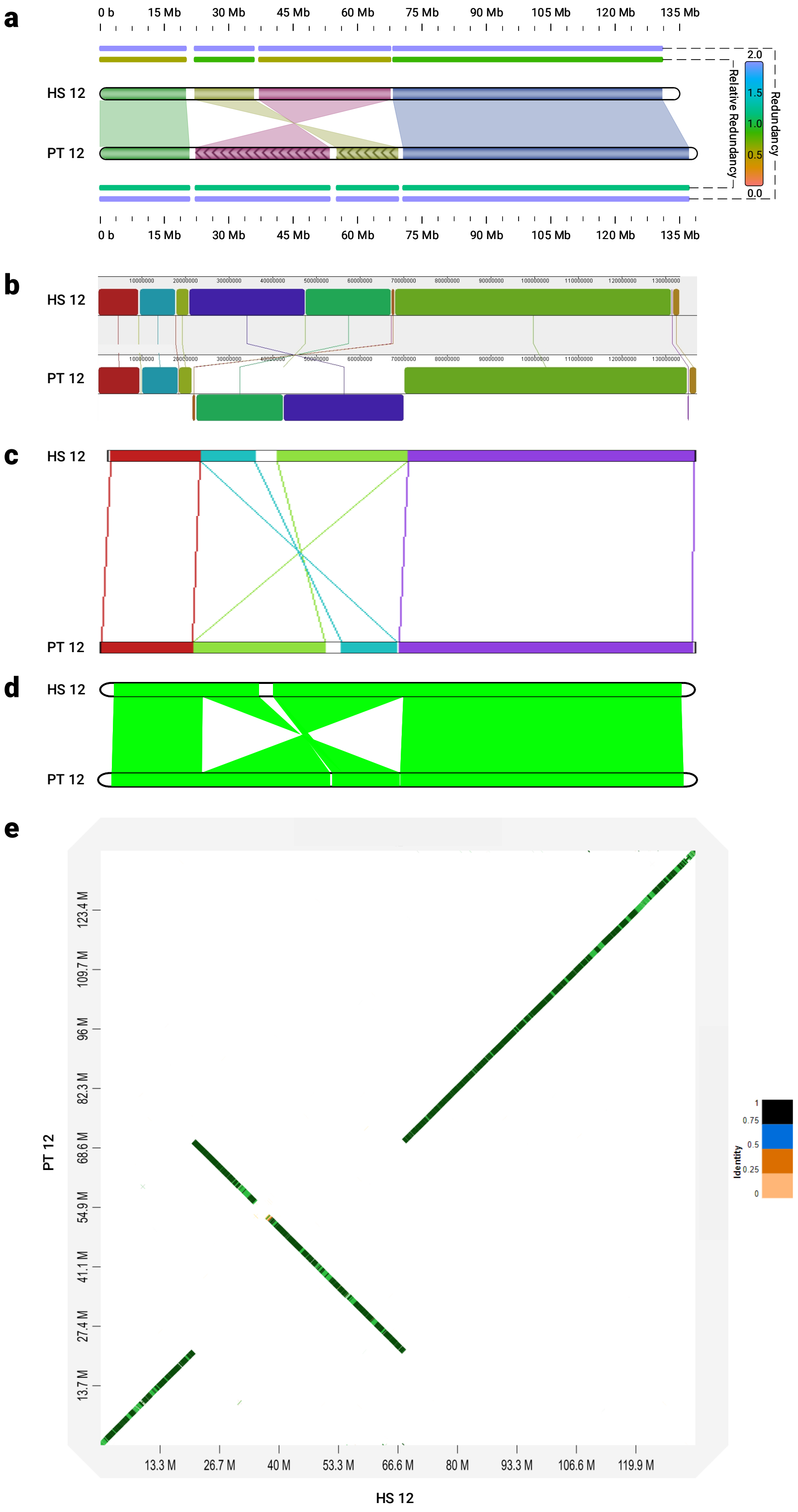


**Figure S3.** Comparison of *H. sapiens* chromosome 12 and *P. troglodytes* chromosome 12. (a) Smash++, with $k = 14$ used by an FCM. The blocks smaller than 100 Kb are discarded; (b) progressiveMauve, with LCB weight of 55186; (c) Cinteny [5], with minimum length of synteny block = 1 Kb, maximum gap between adjacent markers = 5 Mb and minimum number of markers = 1; (d) SynBrowser, with the resolution of 150 Kb; (e) D-Genies [6], in "strong precision" mode.

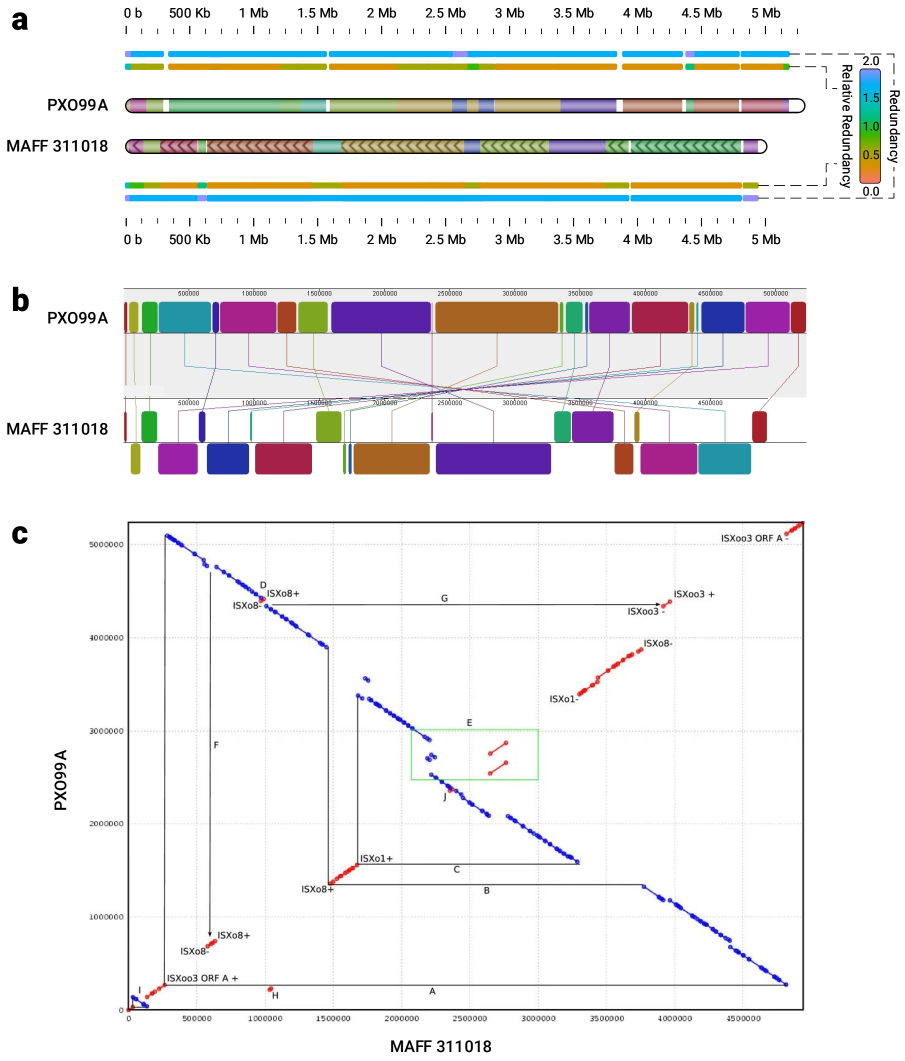


**Figure S4.** Pair-wise comparison of PXO99A and MAFF 311018. (a) Smash++, with k=13 used by an FCM. The blocks smaller than 10 Kb are discarded. In order to make the figure clearer, the shaded paths for connecting corresponding regions are not drawn; (b) progressiveMauve, with LCB weight of 3926; (c) adopted from [7], which employs an alignment-based method to obtain this dot plot. The blue and red colors shows regions of PXO99A that align to the same or opposite strand of MAFF 311018, respectively.

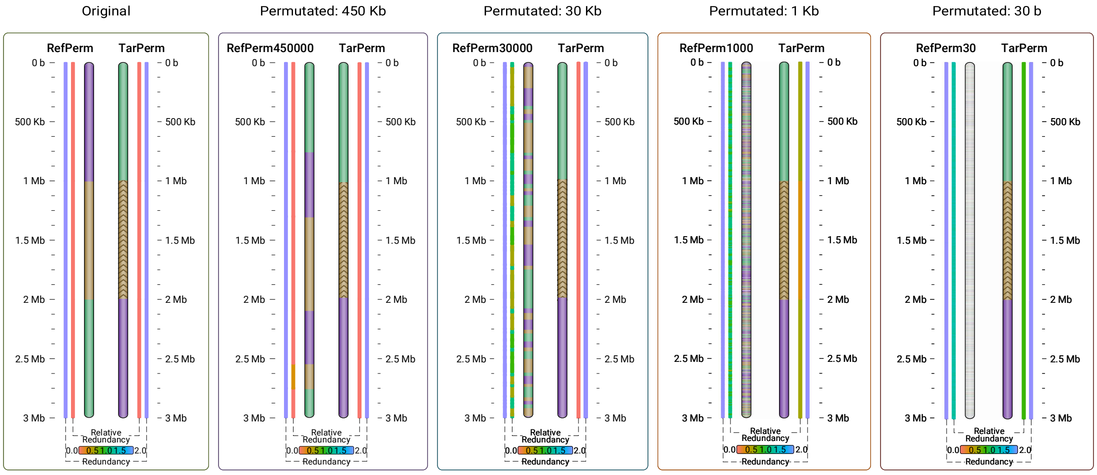


**Figure S5.** Similarity of a target sequence to a reference sequence, when the reference is permutated by blocks of different sizes of 450 Kb, 30 Kb, 1 Kb and 30 b. To run Smash++, an FCM with $k$-mer size of 14 and the threshold of 1.5 was used. It shows the proposed method is resistant to fragmentation of sequence; for example, in the case of 30 b, although the reference is highly fragmented, Smash++ could detect the same three similar regions in the target as the original case.

**Table S1.** Performance of Smash++, in terms of memory and time usage, running on all synthetic and real datasets.

| Dataset (Reference + Target) | Category | Size (b) | Memory (KB) | Elapsed time (s) | User time (s) | System time (s) |
|---|---|---|---|---|---|---|
| Small | Synthetic | 3000 | 1,053,516 | 6.08 | 2.16 | 4.53 |
| Medium | Synthetic | 200,000 | 1,053,392 | 4.63 | 2.08 | 3.15 |
| Large | Synthetic | 10,000,000 | 1,056,220 | 25.71 | 23.02 | 3.35 |
| XLarge | Synthetic | 200,000,000 | 1,102,420 | 616.16 | 604.86 | 9.79 |
| Mutate | Synthetic | 120,000 | 1,053,960 | 2.68 | 1.23 | 1.80 |
| PermOrig | Synthetic | 6,000,000 | 1,055,528 | 22.52 | 17.45 | 5.85 |
| Perm450000 | Synthetic | 6,000,000 | 1,055,324 | 27.27 | 19.36 | 8.66 |
| Perm30000 | Synthetic | 6,000,000 | 1,055,148 | 99.26 | 52.91 | 47.08 |
| Perm1000 | Synthetic | 6,000,000 | 1,054,888 | 965.14 | 549.95 | 415.99 |
| Perm30 | Synthetic | 6,000,000 | 1,057,788 | 16,852.20 | 10,603.13 | 6249.22 |
| GGA14_MGA16 | Real | 31,542,564 | 1,067,628 | 94.79 | 87.73 | 7.75 |
| GGA18_MGA20 | Real | 22,419,393 | 1,063,128 | 79.55 | 72.66 | 7.65 |
| HS12_PT12 | Real | 268,046,531 | 1,135,156 | 617.06 | 604.24 | 9.98 |
| PXO99A_MAFF311018 | Real | 10,324,186 | 1,057,792 | 106.30 | 79.61 | 27.71 |
| CompSynth | Synthetic | 2,000,000 | 1,054,664 | 11.43 | 7.51 | 4.62 |
| CompReal | Real | 2,228,294 | 1,054,596 | 24.06 | 13.64 | 11.18 |

### Note S1 Software manual

Smash++ is implemented in the C++ language and is publicly available at `https://github.com/smortezah/smashpp`, under GNU GPL v3 license. This tool is able to find and visualize rearrangements in a pair of DNA sequences. It is recommended to use as input bare sequences, i.e., without header or quality scores, although, FASTA and FASTQ formats are also supported. In the following sections, we describe installing and running the Smash++ tool.

#### S1.1 Install

To install Smash++ on various operating systems, follow the instructions below. Note that the precompiled executables are available for 64 bit operating systems in the “experiment/bin/” directory.

##### Conda

```
conda install -c cobilab smashpp
```

##### Linux

- Install “git” and “cmake”:

```
sudo apt update
sudo apt install git cmake
```

- Clone Smash++ and install it:

```
git clone https://github.com/smortezah/smashpp.git
cd smashpp
./install.sh
```

##### macOS

- Install “Homebrew”, “git” and “cmake”:

```
/usr/bin/ruby -e "$(curl -fsSL https://raw.githubusercontent.com/Homebrew/
    install/master/install)"
brew install git cmake
```

- Clone Smash++ and install it:

```
git clone https://github.com/smortezah/smashpp.git
cd smashpp
./install.sh
```

##### Windows

After installing “wsl” (Windows Subsystem for Linux), the commands to install Smash++ would be the same as for Linux/macOS:

```
git clone https://github.com/smortezah/smashpp.git
cd smashpp
./install.sh
```

#### S1.2 Run

Various options are provided with the interface of the proposed tool, which are described in Table S2. The commands for running Smash++ have the form of the following:

```
./smashpp [OPTIONS] -r <REF-FILE> -t <TAR-FILE>
```

**Table S2.** Options provided by Smash++ interface.

| Flag | Input | Description |
|---|---|---|
| *Required* | | |
| `-r` | Seq/FASTA/FASTQ | Reference file. Spaces in the name is supported. |
| `-t` | Seq/FASTA/FASTQ | Target file. Spaces in the name is supported. It is recommended to have short names. |
| *Optional* | | |
| `-l` | Integer: $[0, 6]$ Default: 3 | Level of compression. |
| `-m` | Integer: $[1, 2^{32} - 1]$ Default: 50 | Minimum segment size. Only the regions that have greater sizes than this value would be considered for compression. |
| `-e` | Float: $[0.0, 100.0]$ Default: 2.0 | Entropy of 'N' bases. In implementation of the reference-based compression, we replace 'N's in references and targets with 'A's and 'T's, respectively. On reference-free compression, we replace them with 'A's, in both references and targets. If a user tends to replace 'N' bases in a sequence with a normal distribution of 'A', 'C', 'G' and 'T's, he/she can employ the GOOSE toolkit `https://github.com/pratas/goose`. |
| `-n` | Integer: $[1, 255]$ Default: 4 | Number of threads. Creating multiple finite-context models and substitution-tolerant Markov models can be done in a multi-threaded fashion. |
| `-f` | Integer: $[1, 2^{32} - 1]$ Default: 256 | Filter size. In the process of finding similar regions in the reference and the target sequences, the information content that would be obtained by compression needs to be filtered. |
| `-ft` | Integer or String: {0/rectangular, 1/hamming, 2/hann, 3/blackman, 4/triangular, 5/welch, 6/sine, 7/nuttall} Default: hann | Filter type (windowing function). Besides Hann window that is used by default to smooth the information content (profile), we have implemented several other windowing functions: Blackman [8], Hamming [9], Nuttall [10], rectangular [11], sine [12], triangular [13] and Welch [14] windows. These functions along with Hann function are given by Equation S1 and are plotted in Fig. S6. |
| `-fs` | Char or String: {S/small, M/medium, L/large} | Filter scale. It automatically chooses filter size. If a user does not use this flag, the "-f" flag will handle the filtering size. |
| `-d` | Integer: $[1, 2^{64} - 1]$ | Sampling steps. Instead of considering the whole information content, we can make samples of it. The default value is $\lceil \min(\lvert\text{ref}\rvert, \lvert\text{tar}\rvert) / 5000 \rceil$. Therefore, if this flag is not entered by user, a maximum number of 5000 values of the information content will be considered. |
| `-th` | Float: $[0.0, 20.0]$ Default: 1.5 | Threshold for entropy. This option can be used for segmenting the filtered information content. |
| `-rb` | Integer: $[-2^{15}, 2^{15}-1]$ | Reference beginning guard. |
| `-re` | Integer: $[-2^{15}, 2^{15}-1]$ | Reference ending guard. |
| `-tb` | Integer: $[-2^{15}, 2^{15}-1]$ | Target beginning guard. |
| `-te` | Integer: $[-2^{15}, 2^{15}-1]$ Default: 0 | Target ending guard. Smash++ is capable of finding even very small similar regions in two sequences. However, we have found experimentally that when it is running in a very sensitive mode, there might be some cases in which the size of similar regions in the reference and the target are not balanced. These cases can be handled by "-rb", "-re", "-tb" and "-te" options, by resizing the beginning and ending guards of reference and target regions, respectively. For example, if "-tb 10" is used, the first 10 bases in each target region will be discarded, which results in smaller regions. Note that when the guard sizes of target regions are increased, the models built from these regions would be slightly different than the original models; consequently, the sizes of reference regions that are detected as being similar to the ones from the target would be altered. Therefore, changing the guard sizes of target regions will affect the sizes of reference regions. The same thing happens for the sizes of target regions, when the guard sizes of reference regions are changed. |
| `-ar` | N/A | Consider asymmetric regions. It makes Smash++ not to balance the similar reference and target regions. |

| | | |
|---|---|---|
| `-nr` N/A | | Do not compute self complexity. It makes the tool not to perform the reference-free compression (self-complexity computation). |
| `-sb` N/A | | Save sequence (input: FASTA/FASTQ). Smash++ accepts as input FASTA and FASTQ files, in addition to bare sequences. In these cases, the input files are first converted to sequences and then processed further. It is possible to save these sequences by this option. |
| `-sp` N/A | | Save profile, *.prf. When the information profile is obtained, Smash++ smoothens then removes it, by default; however, it can be preserved by this option. |
| `-sf` N/A | | Save filtered file, *.fil. The filtered profile is segmented then removed, by default; however, it can be preserved by this option. |
| `-ss` N/A | | Save segmented files, *.$s_i$. |
| `-sa` N/A | | Save all generated files, including profile, filtered and segmented files. |
| `-rm`<br>`-tm` | $k$,[$w$,$d$,]ir,$\alpha$,$\gamma$/$t$,ir,$\alpha$,$\gamma$:...<br>$k$,[$w$,$d$,]ir,$\alpha$,$\gamma$/$t$,ir,$\alpha$,$\gamma$:...<br>Default:<br>$14, 0, 0.001, 0.95/5, 0, 0.001, 0.95$ | Parameters of reference models.<br>Parameters of target models:<br>• $k$ (integer $> 1$): context size,<br>• $w$ (integer $< 2^{64} - 1$): Count-Min-Log sketch width in $\log_2$ form, e.g., set 10 for $w = 2^{10} = 1024$,<br>• $d$ (integer $> 0$): Count-Min-Log sketch depth,<br>• ir (integer: $\{0, 1, 2\}$): inverted repeat, including 0: regular (not inverted), 1: inverted solely, and 2: both regular and inverted,<br>• $\alpha$ (float $> 0$): estimator,<br>• $\gamma$ (float: $[0.0, 1.0)$): forgetting factor,<br>• $t$ (integer $> 0$): threshold (number of substitutions).<br>It is recommended for compression to use "-l" option, since it configures the models automatically. However, using "-rm" and "-tm", the user would be able to manually configure the reference model, for reference-based compression, and the target model, for reference-free compression, respectively. Parameters of models are described in detail in the section "Methods" of the main paper. |
| `-ll` N/A | | List of compression levels. It shows the list of parameters that would be chosen automatically for each model. |
| `-h` N/A | | Usage guide. |
| `-v` N/A | | More information (verbose). |
| `--version` | | Show the version. |

$$w[n] = 1, \quad 0 \le n \le N, \qquad \text{(rectangular)}$$
$$w[n] = 0.54348 - 0.45652\ \cos\left(\tfrac{2\pi n}{N}\right), \quad 0 \le n \le N \qquad \text{(Hamming)}$$
$$w[n] = \sin^2\left(\tfrac{\pi n}{N}\right), \quad 0 \le n \le N \qquad \text{(Hann)}$$
$$w[n] = 0.42659 - 0.49656\ \cos\left(\tfrac{2\pi n}{N}\right) + 0.07685\ \cos\left(\tfrac{4\pi n}{N}\right), \quad 0 \le n \le N \qquad \text{(Blackman)}$$
$$w[n] = 1 - \left|\tfrac{n-N/2}{N/2}\right|, \quad 0 \le n \le N \qquad \text{(triangular/Bartlett)}$$
$$w[n] = 1 - \left(\tfrac{n-N/2}{N/2}\right)^2, \quad 0 \le n \le N \qquad \text{(Welch)}$$
$$w[n] = \sin\left(\tfrac{\pi n}{N}\right), \quad 0 \le n \le N \qquad \text{(sine)}$$
$$w[n] = 0.35577 - 0.48740\ \cos\left(\tfrac{2\pi n}{N}\right) + 0.14423\ \cos\left(\tfrac{4\pi n}{N}\right) - 0.01260\ \cos\left(\tfrac{6\pi n}{N}\right), \quad 0 \le n \le N \qquad \text{(Nuttall)}$$

**(Eq. S1)**

By running Smash++, positions of similar regions in reference and target sequences, relative redundancy and also redundancy (complexity) of the regions is saved in a ".pos" file. This tab-delimited file has a header including:

1. The string "##SMASH++" as a specifier for the Smash++ tool,

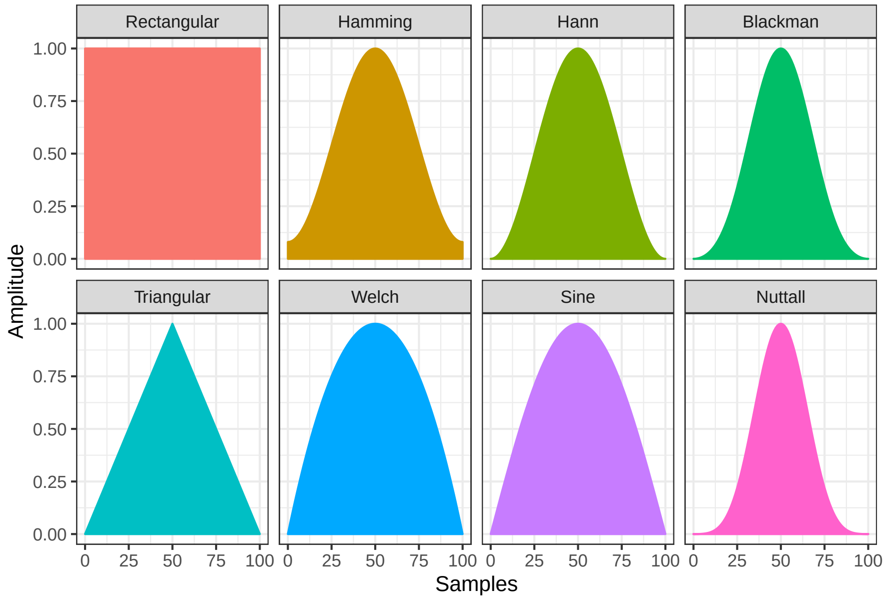


**Figure S6.** Various windowing functions embedded in Smash++.

2. The "PARAM" line to list the parameters used to generate the position file,
3. The "INFO" line to provide the names and sizes of reference and target sequences,
4. The line with the names of columns of the body,

and a body including:

1. RBeg (Reference Begin): beginning of a reference region,
2. REnd (Reference End): end of a reference region,
3. RRelRdn (Reference Relative Redundancy): relative redundancy obtained by compressing the associated target block considering as reference this reference block,
4. RRdn (Reference Redundancy): redundancy (Complexity) of the detected reference block, calculated by reference-free compression,
5. TBeg (Target Begin): beginning of a target region,
6. TEnd (Target End): end of a target region,
7. TRelRdn (Target Relative Redundancy): relative redundancy obtained by compressing the associated reference block considering as reference this target block,
8. TRdn (Target Redundancy): redundancy (Complexity) of the detected target block, calculated by reference-free compression,
9. Inv (Invereted repeat): if the corresponding line is an inverted repeat. "F" (False) means it is regular and "T" (True) means it is inverted.

As an example, the header of the following ".pos" file shows that Smash++ was run by the parameters "-r dataset/REF -t dataset/TAR -l 0 -m 1000" and also the reference "REF" and the target "TAR" are 50,000 base long. The body shows that there is a reference block, from the position 5000 up to 10000, similar to a target block, from the position 2000 to 7000. Relative redundancy of compressing the TAR block, using the REF block as reference, is 1.2473. This number is 1.0187 when the REF block is compressed based on the model of the TAR block. Also, redundancies (complexities) of the REF and the TAR blocks are 1.9010 and 1.8580, respectively. The "F" in the last column shows that this similarity is not of the form inverted repeat. The second record of the body shows an inverted repeat rearrangement.

```
##SMASH++
##PARAM=<-r dataset/REF -t dataset/TAR -l 1 -m 1000>
##INFO=<Ref=REF,RefSize=50000,Tar=TAR,TarSize=50000>
#RBeg  REnd   RRelRdn  RRdn    TBeg   TEnd   TRelRdn  TRdn    Inv
5000   10000  1.0187   1.9010  2000   7000   1.2473   1.8580  F
20000  30000  1.2367   1.9777  40000  30000  1.2545   1.9888  T
```

#### S1.3 Run visualizer

The position file obtained by Smash++ can be visualized by

```
./smashpp -viz
```

The visualizer provides various options that are described in Table S3. The commands for running the Smash++ visualizer are of the form

```
./smashpp -viz [OPTIONS]  -o <SVG-FILE>  <POS-FILE>
```

**Table S3.** Options provided by the Smash++ visualizer interface.

| Flag | Input | Description |
|---|---|---|
| *Required* | | |
| | *.pos file | Position file, generated by the Smash++ tool. Spaces in the name is supported. It can be redirected to Smash++ by standard input ("stdin"). |
| *Optional* | | |
| `-o` | *.svg file<br>Default: map.svg | Output image name. |
| `-rn` | String | Reference name shown on output. |
| `-tn` | String<br>Default: names in position file's header | Target name shown on output.<br>If it has some spaces, use double quotes, e.g. "Seq label". |
| `-l` | Integer: $[1, 6]$<br>Default: 1 | Type of the link between similar regions. |
| `-c` | Integer: $[0, 1]$<br>Default: 0 | Color mode. |
| `-p` | Float: $[0.0, 1.0]$<br>Default: 0.9 | Opacity. |
| `-w` | Integer: $[8, 100]$<br>Default: 10 | Width of the sequence. |
| `-s` | Integer: $[5, 200]$<br>Default: 40 | Space between sequences. |
| `-tc` | Integer: $[1, 255]$ | Total number of colors in the map, which is automatically chosen by default. |
| `-rt` | Integer: $[1, 2^{32}-1]$ | Reference tick size. |
| `-tt` | Integer: $[1, 2^{32}-1]$ | Target tick size. |
| `-th` | Integer: $\{0, 1\}$<br>Default: 1 | Tick human readable: 0=false, 1=true. If it is true, the sizes on axes are printed in the format 1K, 2M, etc. Note that here, 1K is equivalent to 1000 and not 1024, and so on. |
| `-m` | Integer: $[1, 2^{32}-1]$<br>Default: 1 | Minimum block size. Only the regions with greater sizes than this value will be illustrated. |
| `-vv` | N/A | Vertical view of the output image. |
| `-nrr` | N/A | Do not show relative redundancy (relative complexity). |
| `-nr` | N/A | Do not show redundancy (complexity).<br>Smash++ performs reference-based and reference-free compressions to calculate the NRC and redundancy, respectively. If a user does not tend to show them, he/she can turn them off by "-nn" and "-nr" triggers. |
| `-ni` | N/A | Do not show inverse maps. |
| `-ng` | N/A | Do not show regular (not inverse) maps. |

| | | |
|---|---|---|
| | | Smash++ considers by default both regular and reverse complement maps in its calculations. |
| -h | N/A | Usage guide. |
| -v | N/A | More information (verbose). |
| --version | | Show version. |

#### S1.4 Example

This section guides step-by-step employing Smash++ to find and visualize rearrangements between two synthetic sequences. Note that the commands can be run on Linux and macOS, however, they are similar in Windows.

First, install Smash++:

```
git clone https://github.com/smortezah/smashpp.git
cd smashpp
./install.sh
```

Then, copy “smashpp” executable file into “example/” directory and go to that directory:

```
cp smashpp example/
cd example/
```

There is a 1000 byte reference sequence, named “ref”, as the following:

```
TCCCGGTCTTTTAGCGGCCAGGGGCCTGGGCTGTATATCGAAAAGTAATATCCCTTTATGCACCGACCGTAATTATGGACAGCAC
ATATACATTATGAGATTTAAAGATCGCGTGGACGACCACGCGGGCTTATAGCCTCACCTGAGGAAGGGGGTGCCTGCGAGGGAGC
TTGAACCTGTAGCCCCAATCTCGAACGACCTGAGGCTTGTGTGGTCAGAGTTGTGACCAGAGCGATCCCGTTGTCAAATCAACCT
AGAGGAGAGGTAAGGGATACGGGTTACATCTCTCCGCTCAGATTGCTCCTATCGGTAGGAAATATCGGGGATAACCCAATACAAA
ACGCTGAACTGTTCATATTTAGCAAGAAGGGGGGACCGAGGAGCTAAATCAGGGACTATGTAAAATTAGAGTTCTAAGGATAGTA
GCACCCGCGCATGGATCGAACTCCGCCTTGGTTGGGTGTGGATGCCATTGGCGTCACCGTGCTGATAGCCAAATCCCTAGTGAAG
GTAACGTGCGGCCAATTAAATAGTAACTGGGCGTAGATTCTAGCGGTGGCCTCATACGGCACCAATGTCCATCCTCCCTGTGCCT
AACACTATAACTCCAATCCCGTGAGTCTACCATCGCCGAGAATCAGACGGGCAACAAACCCAGGGAGCGAGAGCTACGCGGCTTA
TATCACGGATGCTTCCCCACCTAGGGAAATTTGTTCCGCCTAATCCCGTCCTTCGCTGGTCAGGCCCGGTCTAGCGGATTACTTC
GCCGGTAATGCAGTACAGAATAAATGAAATTCATTAAGAGAATGAAAGGTTATCAGCGAAGCCTTAAGGTCCAACAAGACGTCGA
TATTCTGAAGTAAACTGGTTGACATGTGTGTAACATGAAGCACGCGCTTATTGATATTATCCGCAACCCCACGGCTGGCGGGAAT
CAACCGGCGTCCAGTTCGAACAAGACAGTGCGCTACGCATTACGAAATAGGCTTCGTGTTGCTGT
```

and a 1000 byte target sequence, named “tar”, in this directory:

```
CAGCAACACGAAGCCTATTTCGTAATGCGTAGCGCACTGTCTTGTTCGAACTGGACGCCGGTTGATTCCCGCCAGCCGTGGGGTT
GCGGATAATATCAATAAGCGCGTGCTTCATGTTACACACATGTCAACCAGTTTACTTCAGAATATCGACGTCTTGTTGGACCTTA
AGGCTTCGCTGATAACCTTTCATTCTCTTAATGAATTTCATTTATTCTGTACTGCATTACCGGCGAAGTAATCCGCTAGACCGGG
CCTGACCAGCGAAGGACGGGATTAGGCGGAACAAATTTCCCTAGGTGGGGAAGCATCCGTGATATAAGCCGCGTAGCTCTCGCTC
CCTGGGTTTGTTGCCCGTCTGATTCTCGGCGATGGTAGACTCACGGGATTGGAGTTATAGTGTTAGGCACAGGGAGGATGGACAT
TGGTGCCGTATGAGGCCACCGCTAGAATCTACGCCCAGTTACTATTTAATTGGCCGCACGTTACCTTCACTAGGATCCCGGTCTT
TTAGCGGCCAGGGGCCTGGGCTGTATATCGAAAAGTAATATCCCTTTATGCACCGACCGTAATTATGGACAGCACATATACATTA
TGAGATTTAAAGATCGCGTGGACGACCACGCGGGCTTATAGCCTCACCTGAGGAAGGGGGGGCCTGCGAGGGAGCTTGAACCTGT
AGCCCCAATCTCGAACGACCTGAGGCTTGTGTGGTCAGAGTGGTGACCAGAGCGATCCCGTTGTCAAATCAACCTAGAGGAGAGG
TAAGGGATACGGGTTACATCTCTCCGCTCAGATTGCTCCTATCGGTAGGAAATATCGGGGATAACCCAATACAAAACGCTGAACT
GTTCATATTTAGTAAGAACGGGTGACCGAGGAGCTAAATCAGGGACTATGTAAAATTAGAGATCTAAGGATAGTAGCACCCGCGC
ATGGATCGAACTCCGCCATGGTTTGGTGTCGATGCCATTGGCGTCACCGTGCTGATAGCCAAATC
```

Next, run Smash++ and the visualizer:

```
./smashpp -r ref -t tar
./smashpp -viz -o example.svg ref.tar.pos
```

Results are shown in Fig. S7. Note that the position file is saved as “ref.tar.pos” and the output plot is saved as “example.svg”.

**a**

```
##SMASH++
##PARAM=<-r ref -t tar>
##INFO=<Ref=ref,RefSize=1000,Tar=tar,TarSize=1000>
#RBeg   REnd   RRelRdn   RRdn    TBeg   TEnd   TRelRdn   TRdn    Inv
0       510    0.5108    2.0000  497    1000   0.5120    2.0000  F
498     1000   0.1518    2.0000  508    0      0.1551    2.0000  T
```

**b**

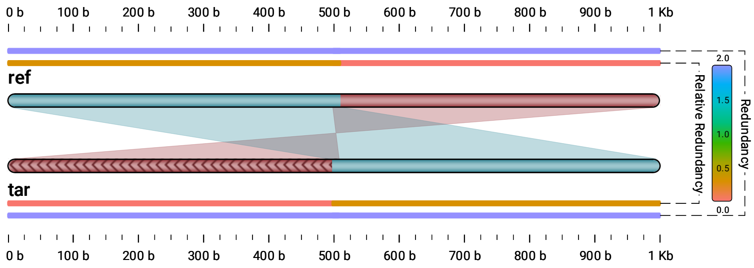


**Figure S7.** An example of running Smash++ on two 1000 base sequences. (a) the position file and (b) output of the visualizer. One similar region in regular mode and another one in inverted mode are detected.
